# Reduced methane emissions in transgenic rice genotypes are associated with altered rhizosphere microbial hydrogen cycling

**DOI:** 10.1101/2024.10.07.617079

**Authors:** Ling-Dong Shi, Maria Florencia Ercoli, Jack Kim, Artur Teixeira de Araujo Junior, Subah Soni, Tracy Satomi Weitz, Alexandra M. Shigenaga, Ilija Dukovski, Rohan Sachdeva, Halbay Turumtay, Katherine B. Louie, Benjamin P. Bowen, Henrik V. Scheller, Daniel Segrè, Trent R. Northen, Pamela C. Ronald, Jillian F. Banfield

## Abstract

Rice paddies contribute substantially to atmospheric methane (CH_4_) and these emissions are expected to increase as the need to feed the human population grows. Here, we show that two independent rice genotypes overexpressing genes for *PLANT PEPTIDES CONTAINING SULFATED TYROSINE* (*PSY*) reduced cumulative CH_4_ emissions by 38% (PSY1) and 58% (PSY2) over the growth period compared with controls. Genome-resolved metatranscriptomic data from rhizosphere soils reveal lower ratios of gene activities for CH_4_ production versus consumption, decrease in activity of H_2_-producing genes, and increase in bacterial H_2_ oxidation pathways in the PSY genotypes. Metabolic modeling using metagenomic and metabolomic data predicts elevated levels of H_2_ oxidation and suppressed H_2_ production in the PSY rhizosphere. The H_2_-oxidizing bacteria have more genes for utilization of gluconeogenic acids than H_2_-producing counterparts, and their activities were likely stimulated by the observed enrichment of gluconeogenic acids (mostly amino acids) in PSY root exudates. Together these results suggest that decreased CH_4_ emission is due to the reduction of H_2_ available for hydrogenotrophic methanogenesis. The combination of rice phenotypic characterization, microbiome multi-omic analysis, and metabolic modeling described here provides a powerful strategy to discover the mechanisms by which specific plant genotypes can alter biogeochemical cycles to reduce CH_4_ emissions.

## Introduction

Methane (CH_4_) is a greenhouse gas with a 27-30 times greater warming effect than carbon dioxide (CO_2_) over a 100-year period. Atmospheric CH_4_ has recently increased to a record-high concentration of 1932.24 ppb (November, 2023)^1^. One of the most important CH_4_ sources is the waterlogged, anaerobic soils of rice paddies, as they support the growth of methanogenic archaea which catalyze the final step in anaerobic decomposition of organic matter to generate CH_4_^2,3^. Although produced CH_4_ can be partially oxidized *in situ* through aerobic and/or anaerobic methane oxidation^4,5^, the imbalance between CH_4_ production and oxidation leads to rice paddies contributing 7-17% of global CH_4_ emissions^6,7^. Emissions may be further exacerbated by the expansion and intensification of rice cultivation to feed the fast-growing human population^8^, and by global warming resulting from anthropogenic climate change^9^.

Numerous approaches have been explored to mitigate CH_4_ emissions from rice paddies, primarily focusing on farming practices such as managing organic matter, water, and nitrogen^9^. Genetic optimization of rice plants, for example, to allocate less photosynthate to roots, is an alternative CH_4_ mitigation strategy. Transgenic introduction of a single transcription factor gene, barley *SUSIBA2*, into rice plants confers more nutrients to aboveground biomass and less to roots, therefore reducing CH_4_ formation^10^. A natural loss-of-function allele of *gs3* (a gene on chromosome 3 controlling grain size) in rice has a similar physiological modulation and causes lower production of CH_4_^11^.

Another approach to reduce CH_4_ emissions is to alter the rice root phenotype. Through programmed cell death, rice roots develop aerenchyma, which facilitates gas diffusion, while simultaneously, the deposit of suberin and lignin forms complete apoplastic barriers in the outer root layers that reduce radial oxygen loss^12,13^. It has been reported that rice genotypes with a higher capacity for internal oxygen diffusion grow larger root systems in waterlogged soils and are associated with lower CH_4_ emissions due to their stronger ability to oxygenate the soil^14,15^. Previously, we and others have shown that overexpression of *PLANT PEPTIDES CONTAINING SULFATED TYROSINE* (*PSY*) genes in *Arabidopsis thaliana* promotes root growth by controlling cell elongation, triggering specific signaling pathways and activating genes involved in the production of secondary metabolites that can be secreted as part of root exudates^16–19^. Given that soil microorganisms inhabit the rice root rhizosphere and that substrates for CH_4_ production (e.g., H_2_/CO_2_ and acetate) by methanogenic archaea ultimately derive from root exudates^20^, we hypothesized that modulation of rice root phenotype would affect microbiome activity and CH_4_ cycling. To this end, we generated rice plants with altered seminal roots by overexpressing the rice *PSY* genes (*OsPSY1* and *OsPSY2*) and cultivated them, along with the control Kitaake, using agronomic soil in a greenhouse. Plant physiological traits including root exudate chemistry, CH_4_ emissions, rhizosphere microbial community composition and *in situ* activities, were compared over the rice growth period.

## Results

### Generation of rice plants overexpressing rice *PSY1* and *PSY2* genes

To identify putative rice *PSY* genes, we surveyed the Nipponbare and Kitaake genomes using bioinformatic approaches as previously described^17,21^, and identified nine candidate *PSY* genes (Supplementary Fig. 1a). Phylogenetic analysis revealed that the active peptide domain sequence is highly conserved across Arabidopsis (AtPSYs) and rice (OsPSYs) (Supplementary Fig. 1b), suggesting that OsPSY peptides may regulate rice root growth similar to AtPSY peptides^16,17,19,22,23^ (Supplementary Fig. 1b). Supporting this hypothesis, we demonstrated that Kitaake seedlings grown on solid media or hydroponically developed longer seminal roots when treated with synthetic OsPSY1 (Fig. 1a and b). Publicly available transcriptomic datasets^24,25^ reveal that *OsPSY1*, also known as *CHALKY GRAIN 5* (*OsCG5*)^26^, is predominantly expressed in spikelets, with lower expression levels in other rice tissues including roots (Supplementary Fig. 1c and e). We hypothesize that the root growth effect observed after synthetic peptide treatment may be due to the expression of leucine-rich repeat receptor-like kinases (LRR-RLKs) in root tissues that can recognize OsPSY1, similar to the abundance of LRR-RLKs observed in Arabidopsis that can bind PSY peptides^23^. In contrast, other members of this gene family, such as *OsPSY2*, displayed more ubiquitous expression across different rice organs (Supplementary Fig. 1d). Specifically, *OsPSY2* expression in rice roots increased progressively in the differentiation zone (DZ), a pattern previously noted for *AtPSY1* promoter activity^19^ (Supplementary Fig. 1e).

**Fig. 1.**
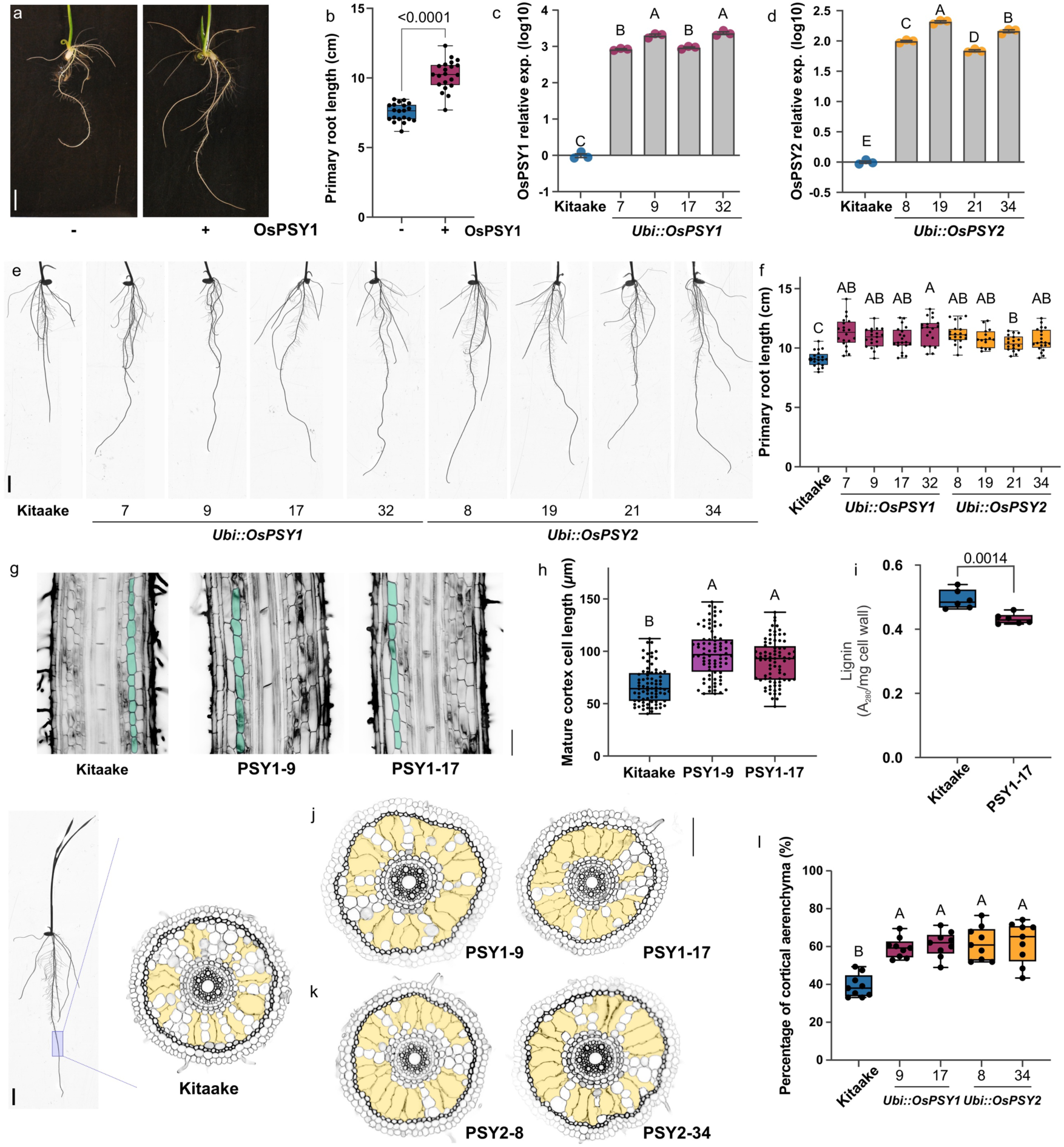
Overexpression of OsPSY1 and OsPSY2 enhanced primary root growth in rice. (**a**) Root phenotype and (**b**) root growth in Kitaake seedlings grown hydroponically for 7 days on 1x MS with or without 500 nM of synthetic OsPSY1 (n=20 seedlings). Expression of (**c**) *OsPSY1* or (**d**) *OsPSY2* in four independent homozygous transgenic lines derived from parental lines *Ubi::OsPSY1* (#7, #9, #17, and #32) and *Ubi::OsPSY2* (#8, #19, #21, and #34), compared with the Kitaake control. Expression was measured by RT-qPCR in three biological replicates and normalized to the mean value obtained in control plants. (**e**) Root phenotype and (**f**) root growth in seedlings of Kitaake, and independent homozygous transgenic lines *Ubi::OsPSY1*(#7, #9, #17, and #32) and *Ubi::OsPSY2* (#8, #19, #21, and #34) grown for 7 days on 1X MS vertical plates (n=20 seedlings). (**g**, **h**) Quantification of cortical cell length (highlighted in green) in the differentiation zone of Kitaake and *Ubi::OsPSY1* (#9 and #17) seedlings grown for 7 days on 1x MS vertical plates (n=4 seedlings). (**i**) lignin content (6 replicates of 8 complete seedlings root system). (**j-l**) Cross-sections depicting the percentage of aerenchyma formation (highlighted in yellow) at 2.5-3.5cm from the root tip in Kitaake, *Ubi::OsPSY1* (#9 and #17) (**j**) and *Ubi::OsPSY2* (#8 and #34) (**k**) independent homozygous transgenic seedlings grown for 7 days on 1x MS vertical plates (n=9 seedlings). On the left, the blue shaded box in the rice seedling indicates the region from which root tissue was collected for imaging (scale bar is 1 cm). In (**b**), the data shown are box and whisker plots combined with scatter plots; each dot indicates the measurement of the designated parameter listed on the y-axis of the plot. P values are calculated by two-tailed Student’s t-test. In (**f**), (**h**), and (**k**), the data shown are box and whisker plots combined with scatter plots; each dot indicates the measurement of the designated parameter listed on the y-axis of the plot. Different letters indicate significant differences, as determined by one-way ANOVA followed by Tukey’s multiple comparison test (P < 0.05). Scale bars in (**a**), (**e**), are 1 cm. Scales bar in (**g**), (**j**) and (**k**) are 100 μm.

To further investigate the roles of *OsPSY1* and *OsPSY2* in rice development, we independently overexpressed each gene in the Kitaake genetic background. We generated more than 50 independently transformed lines using the constitutive UBIQUITIN (Ubi) promoter to drive the expression of *OsPSY1* and *OsPSY2* (*Ubi::OsPSY1* and *Ubi::OsPSY2*). We self-pollinated the resulting transgenic plants and conducted analyses on the progeny derived from four independently transgenic lines, including *Ubi::OsPSY1#7*, *Ubi::OsPSY1#9* (hereafter referred to as PSY1-9), *Ubi::OsPSY1#17* (PSY1-17), *Ubi::OsPSY1#32*, *Ubi::OsPSY2#8* (PSY2-8), *Ubi::OsPSY2#19*, *Ubi::OsPSY2#21* and *Ubi::OsPSY2#34* (PSY2-34). Expression analysis in root tissue showed that progeny derived from the *Ubi::OsPSY1* and *Ubi::OsPSY2* transgenic lines accumulated significantly higher levels of *OsPSY1* or *OsPSY2* transcripts compared with Kitaake (Fig. 1c and d). Consistent with the synthetic peptide treatment analysis, rice plants expressing higher levels of *OsPSY1* or *OsPSY2* exhibited longer seminal roots (Fig. 1e and f) and also carried the transgene (Supplementary Fig. 1f and g). Given that in Arabidopsis, *AtPSY1* regulates root growth by controlling the extent to which cells elongate before reaching their final differentiated size^19,23^, we measured the length of mature cortical cells in the DZ of PSY1-9 and PSY1-17 genotypes. Consistent with the phenotypes observed in Arabidopsis, we found that the roots of rice genotypes accumulating higher levels of *OsPSY1* exhibited longer mature cortical cells (Fig. 1g and h). Additionally, the amount of lignin in PSY1-17 plants was significantly lower compared with Kitaake (Fig. 1i). A similar phenotype was observed in the roots of Arabidopsis plants with higher levels of *AtPSY1,* where lignin deposition in the tracheary elements of the protoxylem occurred further away from the root tip compared with the wild type^19^. These results collectively suggest that PSY peptides from different species may regulate root development through similar molecular mechanisms. Finally, we assessed aerenchyma formation in a selected subset of the rice PSY1 (PSY1-9 and PSY1-17) and PSY2 (PSY2-8 and PSY2-34) genotypes. We found that in all the transgenic lines tested, the percentage of aerenchyma formation was significantly higher compared with Kitaake. However, no differences were observed between the PSY1 and PSY2 lines (Fig. 1j-l). Given the observed phenotypic modifications in roots of rice seedling, particularly the longer seminal roots and enhanced aerenchyma formation, we hypothesized that PSY genotypes may impact the rhizosphere microbial community and associated CH_4_ cycling.

### Growth of PSY rice genotypes in agronomic soils mitigated CH_4_ emissions compared with controls

To assess the effect of PSY overexpression on CH_4_ emissions, we cultivated three independent transgenic rice lines — PSY1-9, PSY1-17, and PSY2-34 — along with the control Kitaake plants in a greenhouse using soils collected from a rice paddy in Davis, CA. The cultivations were conducted in two separate experiments: EXP1 included PSY1-17, PSY1-9 and Kitaake, and EXP2 included PSY1-17, PSY2-34 and Kitaake. Both experiments included bulk soil controls in pots that remained unplanted. Throughout the experiments, all rice lines developed similarly to Kitaake, with heading observed at 55 days after being transferred to the greenhouse^27^ (Fig. 2a). By 70 days post-transplant, both PSY1 and PSY2 genotypes were slightly shorter than Kitaake (Fig. 2b), but they produced significantly more tillers (Fig. 2c), resulting in comparable dry weights of stems and leaves per plant at the end of the plant life cycle (Fig. 2d). Similar trends were observed in EXP1 with PSY1-9 and PSY1-17 (Supplementary Fig. 2a-b). PSY1 genotypes showed partially filled panicles, fewer seeds, and lower thousand-grain weight (TGW) per plant, compared with Kitaake (Fig. 2e-g and Supplementary Fig. 2c). These phenotypes could be attributed to abnormal panicle development, as we found that the tips of the panicles in PSY1 genotypes often contained smaller and unfilled seeds, a phenotype not seen in PSY2 plants (Fig. 2h). Consistently, no significant differences in seed production occurred in the PSY2 genotypes compared with Kitaake (Fig. 2e-h). We also investigated whether the longer seminal root phenotype observed in young seedlings persisted in older PSY plants. Interestingly, PSY1-17 plants developed longer roots 30 days after transplanting, although their root dry biomass was significantly lower than that of Kitaake plants (Fig. 2k and Supplementary Fig. 2d-e). Similar differences in root dry biomass were observed at the end of the plant life cycle (Fig. 2i). Additionally, lignin content in the roots of adult plants revealed that, consistent with findings in young seedlings, PSY1-17 plants accumulated less lignin than Kitaake (Fig. 2j). Although PSY2 plants also had reduced lignin content, their root biomass remained comparable to that of Kitaake (Fig. 2i-j).

**Fig. 2.**
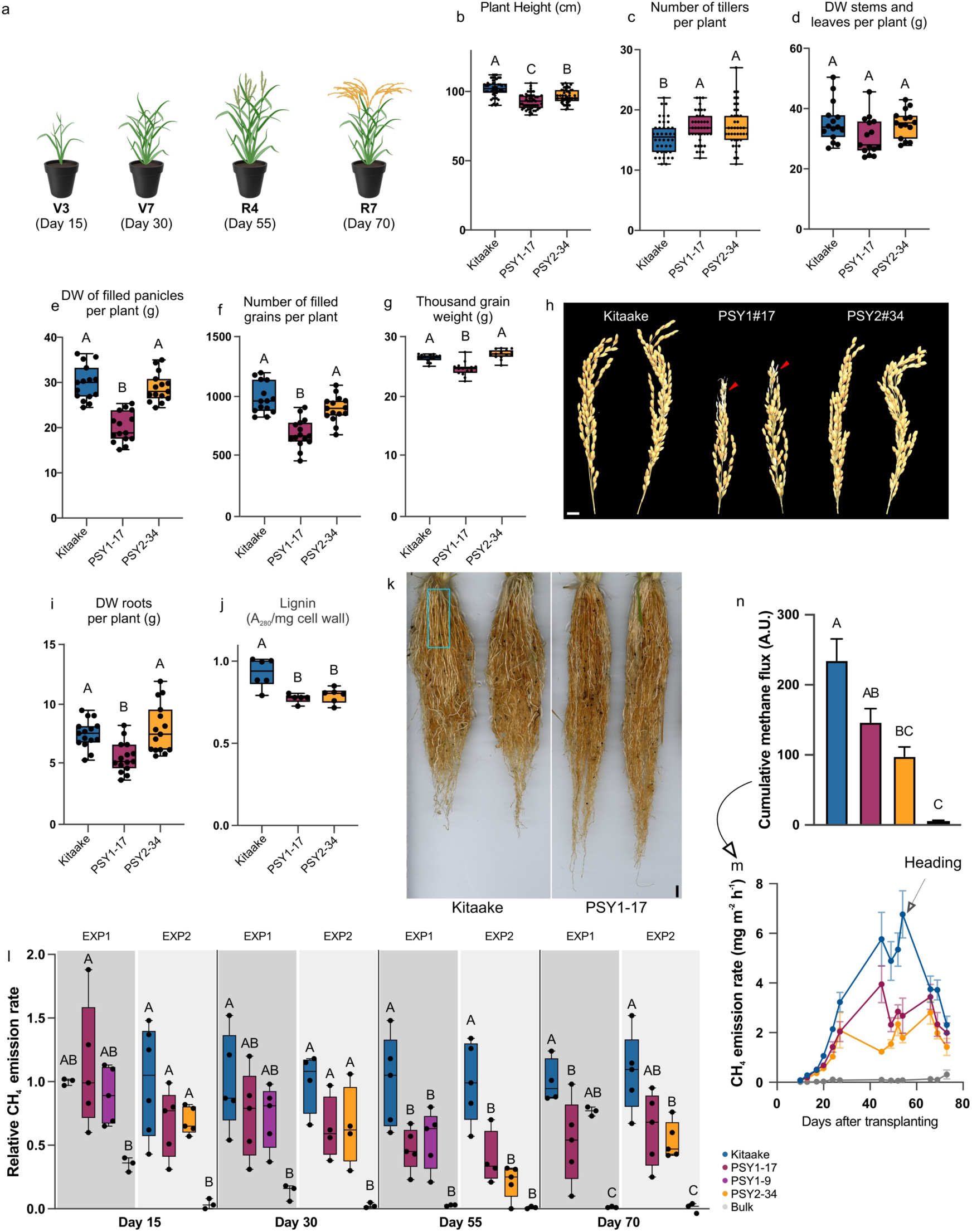
Kitaake plants with elevated expression of *OsPSY1* or *OsPSY2* exhibited reduced CH_4_ emissions. (**a**) Illustrations depicting the plant developmental stage at each time point selected for methane measurements coupled with rhizosphere sample collection in greenhouse EXP1 and EXP2. Phenotypic profiling of Kitaake, PSY1-17, and PSY2-34 grown in EXP2 including (**b**) plant height (cm), (**c**) number of tillers per plant, (**d**) dry weight (DW) from stems and leaves per plant (g), (**e**) DW of filled panicles per plant (g), (**f**) number of filled grain per plant, (**g**) thousand-grain weight (g), (**h**) representative panicles images (scale bar = 1 cm), (**i**) DW from complete root systems per plant, and (**j**) root lignin content. (**k**) Representative image of the root system phenotype from Kitaake and PSY1-17 rice plants 30 days after transplanting in EXP1. Phenotypic measurements were taken 70 days after transplanting for (**b**) (n=50 plants), (**c**) (n=50 plants), and (**j**) (n=6 plants), while (**d**), (**e**), (**f**), (**g**), and (**i**) were analyzed at the end of the plant life cycle 100 days after transplanting (n=15 plants). (**l-n**) For methane emissions in EXP1, we compared Kitaake, PSY1-17 and PSY1-9, whereas in EXP2, Kitaake, PSY1-17 and PSY2-34 were analyzed. In (**l**), the methane emission rates for EXP1 and EXP2 are plotted together with values normalized to the mean value obtained in Kitaake plants for each experiment. (**m**) Methane emission rate plotted over days after transplanting in EXP2. (**n**) Cumulative methane flux was calculated as the area under the curve in (**m**). In (**k**), the light blue square in the Kitaake picture highlights the region of the root system where rhizosphere and lignin samples were collected (scale bar = 1 cm). The data shown in (**b**), (**c**), (**d**), (**e**), (**f**), (**g**), (**i**), (**j**), and (**l**) are box and whisker plots combined with scatter plots; each dot indicates the measurement of the designated parameter listed on the y-axis of the plot. Different letters indicate significant differences, as determined by one-way ANOVA followed by Tukey’s multiple comparison test (P < 0.05).

Despite some phenotypic differences, no significant differences in CH_4_ emissions were observed between the PSY1-9 and PSY1-17 genotypes (EXP1) or between the PSY1 and PSY2 genotypes (EXP2) (Fig. 2l). For these analyses, CH_4_ emissions were measured *in situ* using a Picarro gas analyzer, and soil samples were collected for microbiological analyses throughout the rice growth period (i.e., 15, 30, 55, and 70 days after transplanting; Fig. 2a, k and l). In both experiments, negligible CH_4_ emissions were observed from the bulk soil controls, while differentiated emissions were recorded from pots with various rice plants over the 10-week growth period (Fig. 2l-m). All PSY rice genotypes emitted substantially less CH_4_ compared with Kitaake, with reductions of up to ∼30% observed after heading (55 and 70 days; Fig. 2l). Notably, cumulative CH_4_ emissions throughout the growth period for PSY1 and PSY2 genotypes showed an average decrease of ∼38% and ∼58% relative to Kitaake, respectively (Fig. 2m-n). Given the crucial role of microorganisms in the production and consumption of CH_4_, we hypothesized that distinct microbial communities might accumulate in the rhizospheres of the different rice genotypes, potentially mitigating the rice CH_4_ emissions. Because both PSY1 and PSY2 genotypes displayed comparable reductions in CH_4_ emissions, we focused on PSY1 genotypes for further study.

### Rice PSY genotypes impacted microbial activity but not microbial community structure

To explore the difference in root microbiomes, we extracted and sequenced total DNA and RNA from 72 soil samples (3 replicates for each of the bulk soils and 5 replicates for each planted with the PSY1 rice genotypes and Kitaake control, at each collection time point) in EXP1 (Fig. 2a), generating 974.8 Gbp and 920.7 Gbp of metagenomic and metatranscriptomic raw read data, respectively (Supplementary Table 1). After combining replicates to generate datasets representative of 16 samples, quality trimmed metagenomic reads were assembled and ribosomal protein S3 (rpS3) sequences were used to perform an initial census of microbial diversity, including organism type and abundance^28^. In total, 4,087 rpS3 sequences were identified and grouped into 1,656 non-redundant, approximately species-level clusters.

Phylogenetic analysis based on rpS3 sequences classified all detected microorganisms into at least 4 archaeal and 14 bacterial phyla (Fig. 3a). By mapping metagenomic and metatranscriptomic reads to rpS3 genes, the most abundant and active organisms were Proteobacteria, Chloroflexi, Acidobacteria, Bacteroidetes, Gemmatimonadetes, and Actinobacteria (Fig. 3b and c), consistent with previous soil surveys^29–31^. Between 1 and 5% of sequences were assigned to the “Candidate Phyla Radiation” (CPR) bacteria and DPANN archaea (Diapherotrites, Parvarchaeota, Aenigmarchaeota, Nanoarchaeota and Nanohaloarchaeota), both of which were recently reported in paddy soils^32,33^.

**Fig. 3.**
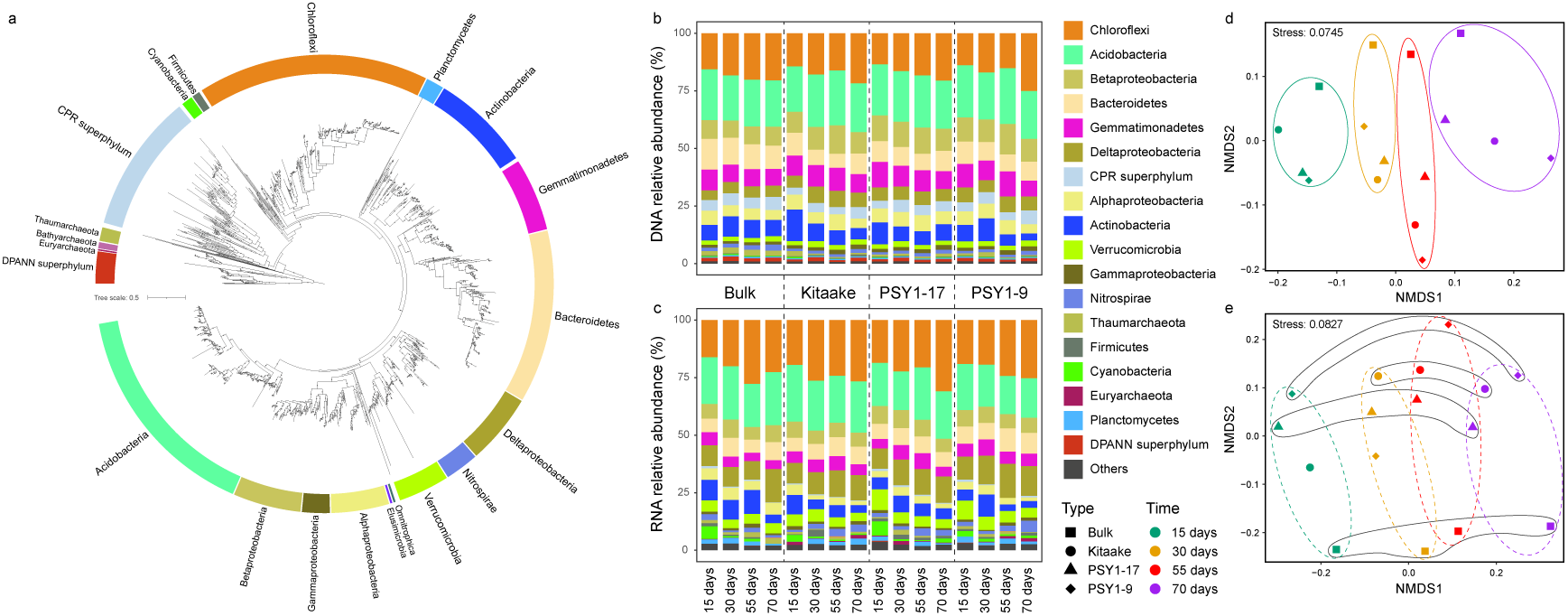
Microbial community structure and activity. (**a**) Microbial community membership based on phylogeny of identified rpS3 amino acid sequences. Black dots indicate support values ≥ 80% calculated based on 1000 replicates. Relative abundance (**b**) and *in situ* activity (**c**) of microbial communities at the phylum level, measured at four time points in the bulk soil and three rice genotypes. “Others” indicates the summed values for rare lineages. Non-metric multidimensional scaling (NMDS) ordination of microbial community structure (**d**) and transcriptional abundance (**e**) with symbol meanings showing in “Type” and “Time” keys. Dissimilarity scores (R) were calculated by analysis of similarity (ANOSIM) test: for community structure (**d**), R=0.539, P=0.001 between rice genotypes (highlighted by ovals), and R=0.0729, P=0.238 between cultivation time; for community activity (**e**), R=0.265, P=0.006 between rice genotypes (highlighted by solid ovals), and R=0.195, P=0.020 between cultivation time (highlighted by dashed ovals).

Non-metric multidimensional scaling analyses (NMDS) show clear differences in community structure and microbial activity between soils collected from the rhizosphere and unplanted bulk soil (Fig. 3d and e), suggesting a strong effect from the rice plants^34^. For the planted groups, cultivation time has a large and statistically significant effect on the community structure (dissimilarity score of up to 0.539) but rice genotype does not (Fig. 3d). Notably, however, microbial activity is significantly influenced by both time and genotype with comparable contributions (dissimilarity scores of 0.265 and 0.195, respectively; Fig. 3e).

### Lower activity of CH_4_-producing genes in microbiomes associated with PSY1 rice genotypes

CH_4_ fluxes from rice paddy soils reflect the combined impact of the activities of methanogens and methanotrophs. A search for methyl-coenzyme M reductases (McrA) that participate in methanogenesis and anaerobic CH_4_ oxidation identified multiple homologs affiliated to *Methanobacterium*, *Methanomassiliicoccus*, *Methanoregula*, *Methanocella*, and *Methanosarcina* (Fig. 4a). No recovered McrA sequences are similar to those found in anaerobic CH_4_-oxidizing archaea, suggesting that anaerobic methanotrophs were either absent or too rare to be detected. Many particulate methane monooxygenases (PmoB) genes that enable aerobic CH_4_ oxidation were identified and taxonomically assigned to several bacterial genera (Fig. 4b). The soluble form of methane monooxygenase was undetectable. These genes are typically only expressed under copper limitation conditions^35^ (e.g., <0.1 μM versus 0.4 μM in our fertilized water; Supplementary Table 2).

**Fig. 4.**
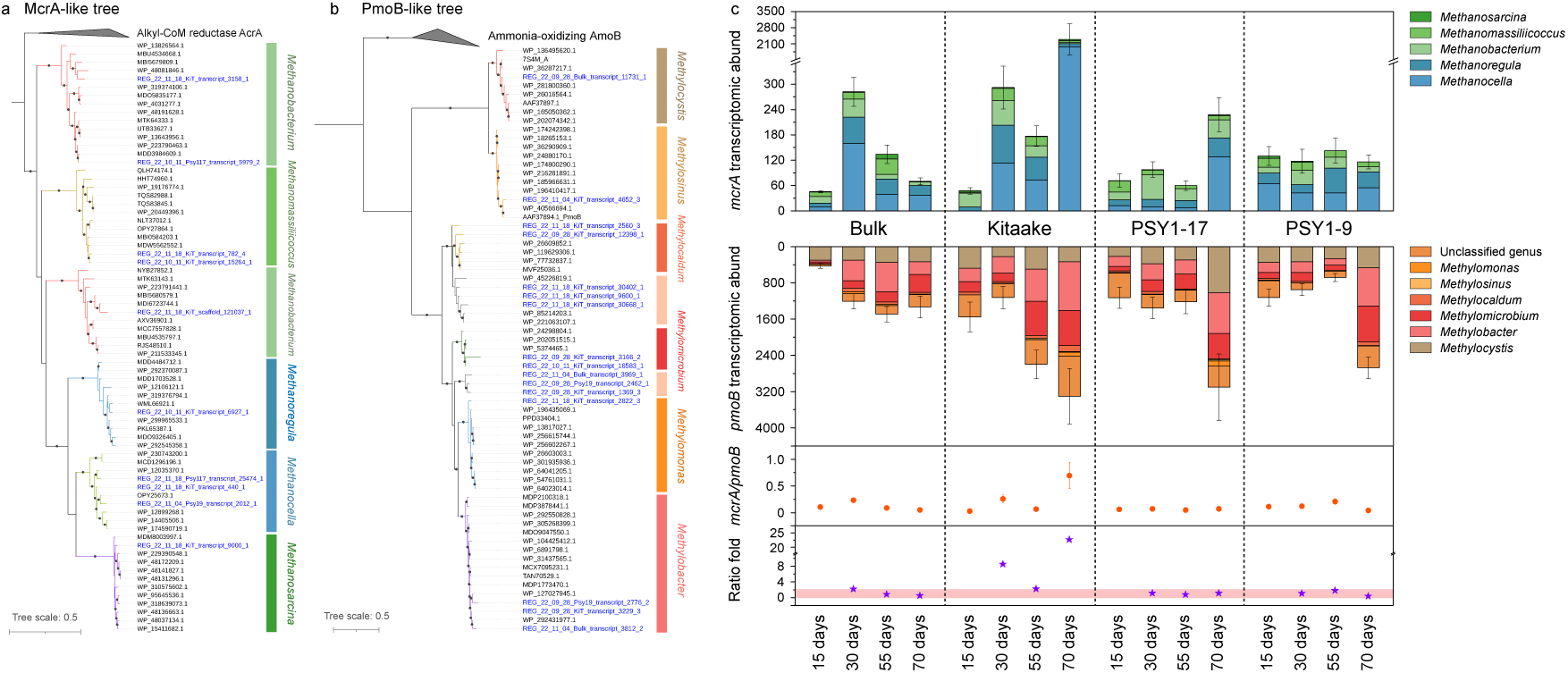
Taxonomic affiliation and *in situ* activity of genes involved in CH_4_ production and oxidation. (**a-b**) Phylogeny of identified McrA and PmoB sequences. Black dots indicate support values ≥ 80% calculated based on 1000 replicates. Blue clades are proteins identified in the study. (**c**) Transcriptional activity and taxonomic affiliation of *mcrA* and *pmoB* genes in the bulk soil and rhizosphere soils of different rice genotypes at four time points during the cultivation period. Error bars indicate standard deviations of sum values in replicates. Ratios of *mcrA* to *pmoB* were calculated by dividing *mcrA* transcriptomic abundances by *pmoB*. Ratio change folds in each group were calculated using ratios at the first sampling point (15 days after transplanting) as baselines. The pink bar indicates the range between 0 and 2.

In general, the transcriptional activity of *mcrA* increased with cultivation time in the three rice genotypes. For example, the activity of *Methanocella mcrA* in the Kitaake genotype increased by approximately one order of magnitude in the last growth stage (70 days, Fig. 3c). The transcriptional activity of *pmoB* also increased with cultivation time, and we observed more active terminal oxidases in microbes associated with the PSY1 genotypes late in the experiment (Supplementary Fig. 3), suggestive of higher O_2_ concentrations. However, the ratio of *mcrA to pmoB* remained stable over the whole growth period, except for in Kitaake (Fig. 4c). Given the importance of the balance between activities that produce and consume CH_4_, we calculated the fold changes of *mcrA*/*pmoB* ratios for each group relative to that at the corresponding first sampling point (i.e., 15 days after transplanting), and observed notable increases in Kitaake but negligible changes in PSY1 genotypes (Fig. 4c). This indicates that, over the rice growth period, *mcrA* activity was promoted much more than *pmoB* in Kitaake, leading to an imbalance between CH_4_ production and consumption, but proportionally in the PSY1 genotypes.

### PSY1 rice genotypes supported higher activities of microbial H_2_-consuming genes but lower activities of H_2_-producing genes

Expressed *mcrA* transcripts were predominantly from *Methanocella*, *Methanoregula*, and *Methanobacterium* (≥90%; Fig. 4c). Archaea of these genera all use H_2_, instead of acetate or methylated compounds, to produce CH_4_^36^. Thus, we identified genes involved in H_2_ production and consumption and tracked their expression levels as a function of genotype and time. Two types of hydrogenases that catalyze the reversible conversion between H_2_ and protons/electrons^37^, namely [NiFe] and [FeFe] hydrogenases, were identified and divided into four and three subtypes, respectively, using protein phylogeny (Fig. 5a and b). The most active clade was [NiFe] Group 1 hydrogenases that oxidize H_2_ (Fig. 5c). The transcriptional activities of this group were generally higher in the two PSY1 genotypes than in the Kitaake control, particularly in the first and last growth stages where the differences were significant (15 and 70 days; Fig. 5d). The [NiFe] Group 4 and [FeFe] Group A1 (further classified by HydDB^38^) are predicted to produce H_2_^39^. Transcript quantification shows [NiFe] Group 4 hydrogenases were more active than [FeFe] Group A1 hydrogenases, and the combined activity of these two H_2_-producing hydrogenases was significantly higher in Kitaake compared with the two PSY1 genotypes, especially in the second and last growth stages (30 and 70 days; Fig. 5d). Given that the microbiomes in the PSY1 genotypes should produce less and consume more H_2_ than in Kitaake, the pool of hydrogen available for CH_4_ production in the soils from the PSY genotypes would be reduced relative to that from the Kitaake control.

**Fig. 5.**
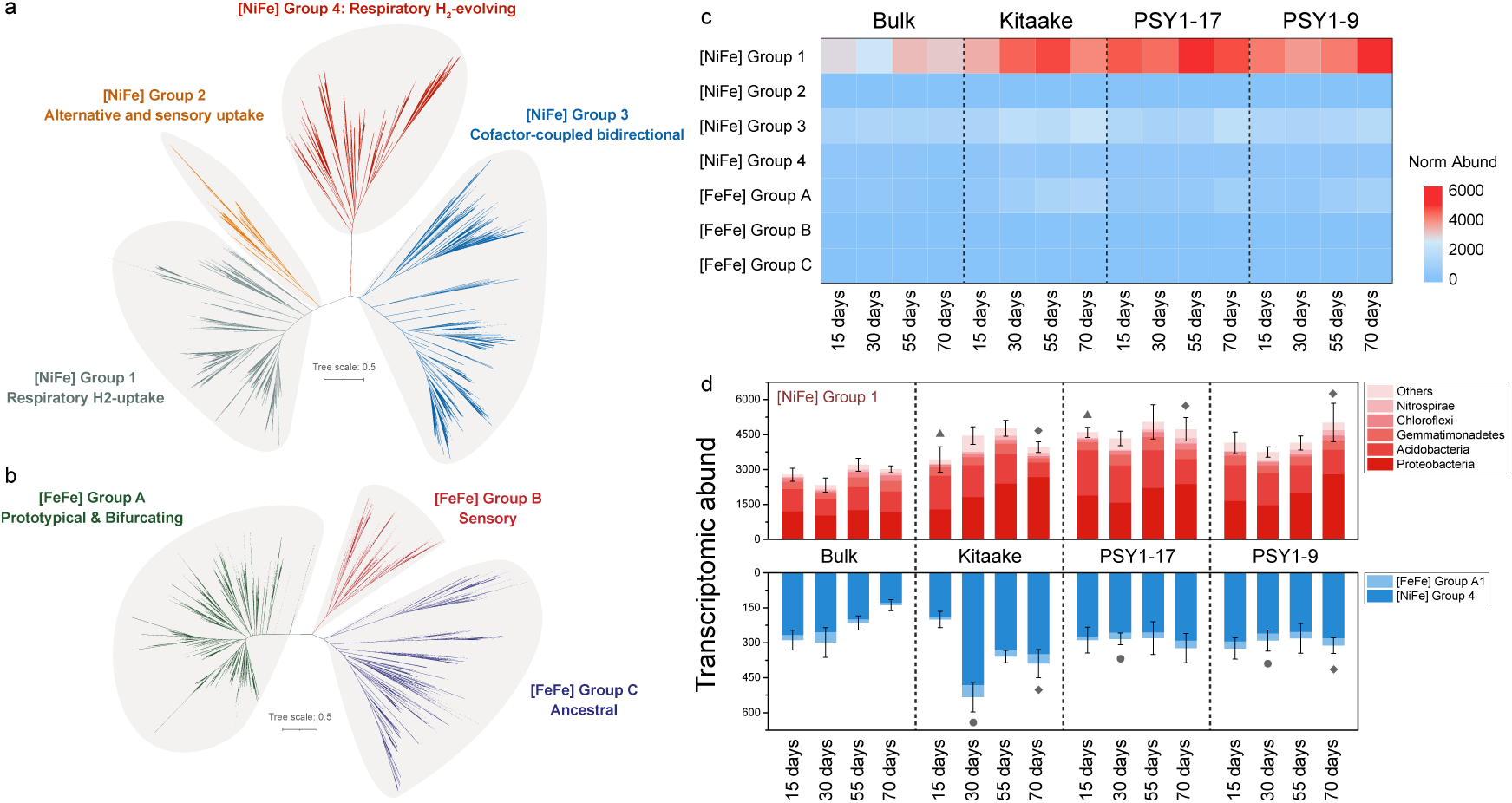
Classification and activity of hydrogenases. Phylogeny of identified [NiFe] hydrogenases (**a**) and [FeFe] hydrogenases (**b**). Black dots indicate support values ≥ 80% calculated based on 1000 replicates. Dashed branches are references. (**c**) Transcriptional activity of classified hydrogenase groups in different rice types over the cultivation period. (**d**) Transcriptomic abundances of [NiFe] Group 1 hydrogenases resolved by organisms of origin, and the combination of [FeFe] Group A1 and [NiFe] Group 4. Error bars indicate standard deviations of sum values in replicates. Symbols (i.e., circles, triangles, diamonds) indicate significant differences between wild type (Kitaake) and PSY genotypes (PSY1-17 and PSY1-9) at the same cultivation time.

An alternative pathway for H_2_ production is as a byproduct of N_2_ fixation (one molecule of H_2_ per N_2_ fixed). The nitrogenase gene (*nifH*) was used as a marker of N_2_ fixation activity and associated H_2_ production^40,41^. The transcriptional activity of *nifH* generally increased with cultivation time and was linked to Delta-, Gamma-, and Alphaproteobacteria (Supplementary Fig. 4). Like H_2_-producing hydrogenases, *nifH* activity was significantly higher in Kitaake than in the PSY1 genotypes, especially in the last two growth stages (55 and 70 days).

### Altered PSY1 root exudates modulate activities of H_2_-metabolizing microorganisms

To explore the influence of rice phenotypes on rhizosphere microbiome activity, we performed metabolomic analysis on root tissues and exudates from axenic rice seedling roots to identify compounds secreted by the PSY1 genotypes. Metabolites were extracted using methanol and profiled on an HPLC-MS platform (see Supplementary Dataset 1). The number of features detected in root tissues or exudates from Kitaake and PSY1 genotypes (including PSY1-9 and PSY1-17 samples) using gradient-based hydrophilic interaction liquid chromatography (HILIC) in positive ionization mode is summarized in Supplementary Fig. 5a and e. Principal component analysis (PCA) demonstrated a separation between PSY1 and Kitaake genotypes, with more distinct sample grouping observed in root tissue features compared to root exudates (Supplementary Fig. 5b and f). While most detected features were common to the transgenic lines and the wild type, some were exclusive to either the PSY1 genotypes or the Kitaake control (Supplementary Fig. 5a and e). Specifically, 2,237 and 2,047 features were shared between Kitaake and PSY1 samples in root tissue and exudates, respectively. Among these, 12% (272 out of 2,237) of the features in PSY1 root tissue samples were differentially abundant compared with wild-type controls (Supplementary Fig. 5g and Supplementary Dataset 2). This percentage was slightly lower in exudate samples, where 8.3% (168 out of 2,047) of features showed differential abundance (Supplementary Fig. 5c and Supplementary Dataset 2). Additionally, several features were either newly detected or depleted in PSY1 genotypes compared with Kitaake. In root tissues, 208 new and 80 depleted features were identified, while exudates exhibited 146 new and 15 depleted features (Supplementary Dataset 2). Overall, more features increased than decreased in PSY1 samples. A subset of these differentially abundant features was putatively identified and classified using Global Natural Products Social Molecular Networking (GNPS) (Supplementary Tables 3-5). Most of the differentially abundant metabolites in PSY1 root tissue and exudate samples belong to the same compound classes, including ‘amino acids’, ‘alkaloids’, and ‘shikimates and phenylpropanoids’ (Supplementary Fig. 5d and h). Similar results were obtained from HILIC in negative ionization mode and reverse-phase (C18) in both positive and negative ionization modes and are detailed in Supplementary Dataset 2, Supplementary Tables 3-5 and Supplementary Figs. 6-8. Altogether, our analysis indicated significant changes in the abundance of compounds in the PSY1 root exudates compared with Kitaake, with most of the metabolites enriched in the PSY1 metabolome classified as gluconeogenic acids (∼90%) (Supplementary Table 6). We also noted the presence of two flavonoids, putatively malvidin-3-glucoside and isorhamnetin 3-rutinoside, that differentially accumulated in exudate samples from the PSY1 genotypes but were absent from Kitaake, a feature observed in previous studies in Arabidopsis^19^.

Next, we analyzed all the microbial genomes reconstructed from the rhizosphere (n=296) and found 17 had transcriptional activities that were significantly up-regulated in soils of rice PSY genotypes relative to the Kitaake control (log2FoldChange ≥ 1, p ≤ 0.05; Supplementary Fig. 9). The genomes of five of the 17 encode H_2_-consuming hydrogenases but none contain H_2_-producing enzymes. This suggests that H_2_-consuming bacteria may respond to the PSY1 rice genotype-derived exudates differently than organisms that produce H_2_. We therefore surveyed the entire H_2_-consuming (n=55) and H_2_-producing (n=36) populations, and calculated glycolytic sugar versus gluconeogenic acid preference values (SAP) using the method described previously^42^. The results show that the H_2_-consuming organisms have much higher preferences for gluconeogenic acids than the H_2_-producing organisms, and their genomes have higher GC contents, likely reflecting this metabolic preference^42^ (Fig. 6a and 6b; Supplementary Table 7). Consequently, H_2_-consuming organisms had higher transcriptional activities in association with the PSY1 genotypes whereas their H_2_-producing counterparts were more active in the Kitaake rhizosphere (Fig. 6c).

**Fig. 6.**
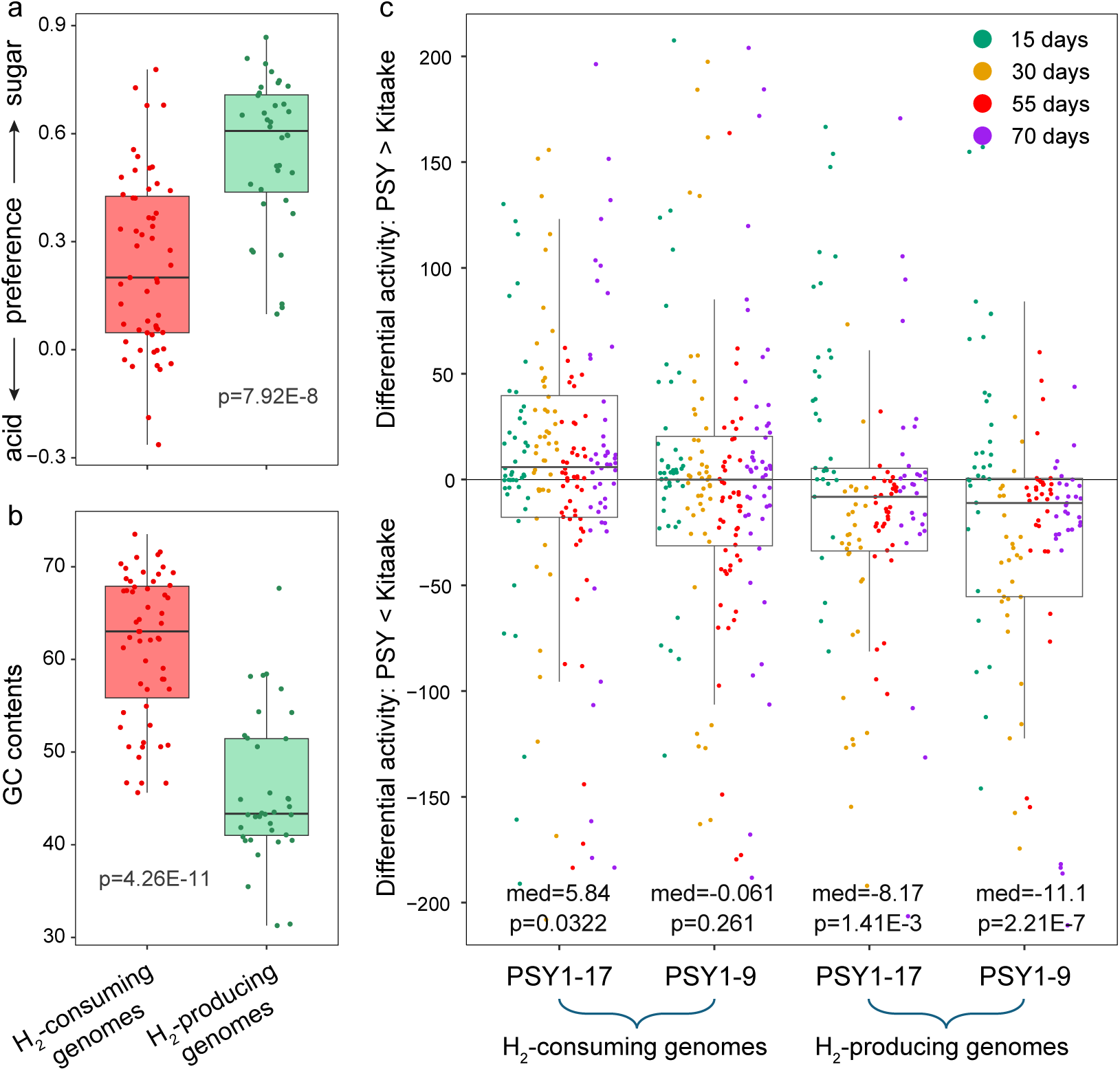
Metabolic preference and activity of H_2_-consuming and H_2_-producing genomes. Predicted sugar-acid-preference values (**a**) and GC contents (**b**) of H_2_-cycling populations. Colored dots indicate genomes in two groups (n=55 and n=36). Box plots show lower and upper quartiles, and median values in each genome group. Whiskers indicate maximum and minimum values excluding outliers. Statistical differences were calculated using the Wilcoxon rank sum test. (**c**) Differential activities of genomes involved in H_2_ cycling. Differential activities were calculated by subtracting overall transcriptional activities in PSY1 rice genotypes from that in Kitaake. Colored dots indicate genomes in different rice growth stages. Box plots show the same information as in (**a**) and (**b**). Median differential activities were statistically compared to zero using one-sample Wilcoxon Signed Rank Test.

To further investigate how rice physiological features affect microbial H_2_ cycling, we built metabolic models using representative, high-quality genomes (completeness >90%, contamination <5%), and predicted H_2_ flux responding to varying PSY1 over-secreted compounds and O_2_ concentrations inferred from aerenchyma densities of the different rice genotypes (Supplementary Table 8). All three modeled H_2_-producing bacteria from different phyla show the same trend, with maximal produced H_2_ decreasing when over-secreted compounds increase from 0.01 to 0.05 fold (Fig. 7a-c). Beyond this range, H_2_ production is predicted to plateau. Microbial growth increased with increasing concentration of over-secreted compounds and plateaued when the H_2_ production plateaued, except for Bacteroidia_37, which continually increased its biomass over the whole simulation (Supplementary Fig. 10a-c). The results suggest that the modeled H_2_-producing bacteria can use over-secreted PSY1 compounds to support growth, but not via pathways that generate H_2_. Increased O_2_ availability has little or no effect on H_2_ production (Fig. 7a-c).

**Fig. 7.**
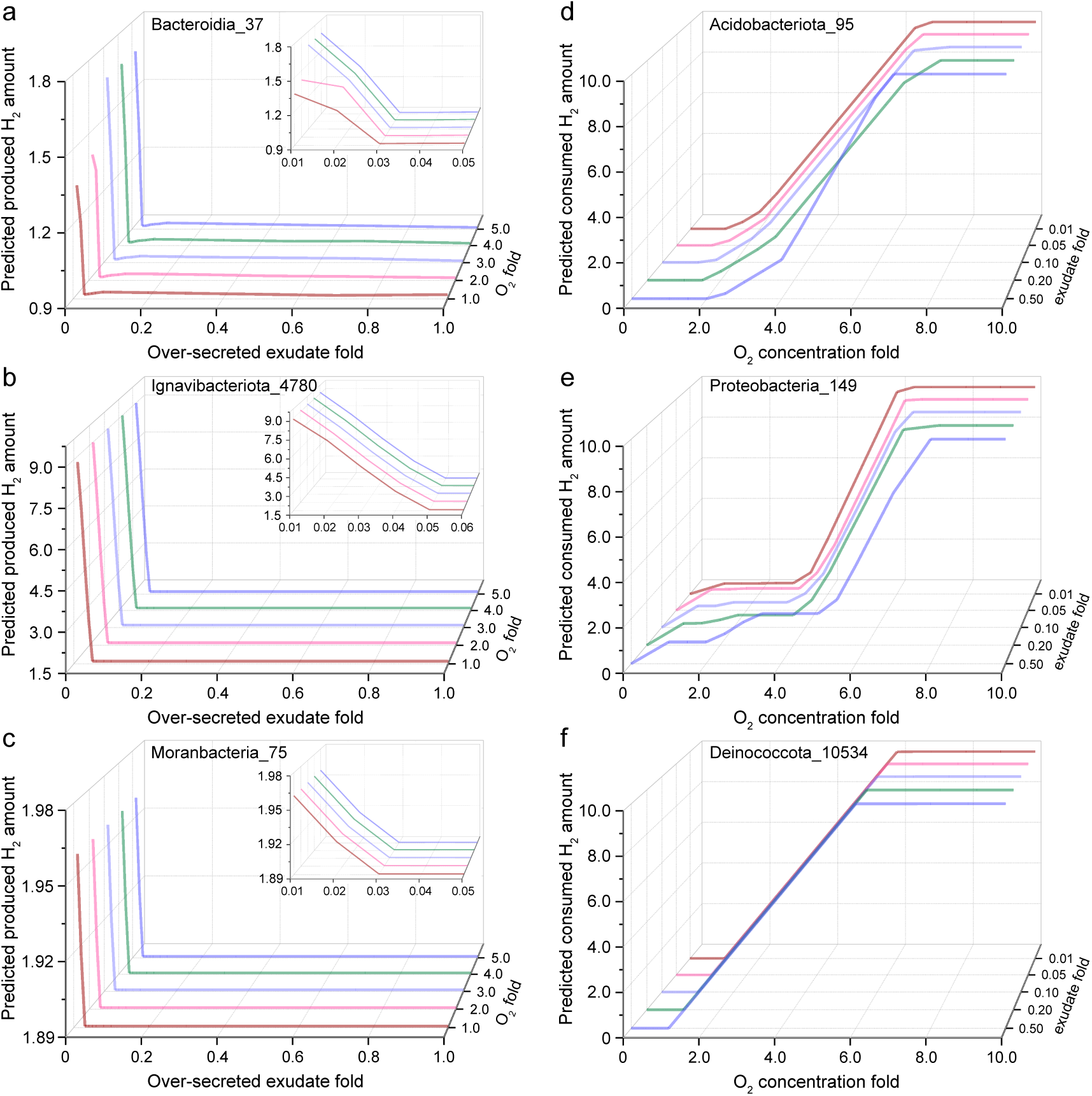
Predicted H_2_ flux by metabolic modeling using representative high-quality genomes. Inferred H_2_ production (**a**-**c**) and H_2_ consumption (**d**-**f**) under varying PSY over-secreted exudate and O_2_ concentrations. Fold changes of over-secreted exudates and O_2_ are defined when compared to essential nutrients that do not differ across rice genotypes (Supplementary Table 8). Insets in panels **a**-**c** show details of H_2_ production at low over-secreted exudate concentrations. H_2_ consumption in panels **d**-**f** was calculated by subtracting final H_2_ concentrations from initial.

The H_2_-consuming organisms are modeled to oxidize more H_2_ (and accordingly yield more biomass) in the presence of higher O_2_ concentrations (Fig. 7d-f and Supplementary Fig. 10d-f). Over-secreted PSY1 compounds have negligible influence on H_2_ consumption, yet also contribute substantially to accumulation of biomass of H_2_-consuming organisms (Supplementary Fig. 10d-e). These modeling results support the metagenomics-based hypothesis that PSY1 rice genotypes enhance the growth and activity of H_2_-consuming organisms due to their distinct exudate profiles and aerenchyma characteristics.

## Discussion

Rice is the most important staple food for half the world’s human population. It is estimated that the global rice production will continue to increase from 506 million tonnes in 2020 to 567 million tonnes by 2030^43^. During cultivation, rice paddies emit millions of tons of CH_4_ annually and contribute ∼10.1% of global agricultural greenhouse gas emissions^44^. Here, we tested whether overexpression of the rice genes, *OsPSY1* or *OsPSY2*, could affect root growth, exudate composition, activities of rhizosphere microbiomes and CH_4_ emissions, potentially providing a route to decrease the climate impact of rice production. To our knowledge, this is the first study to use an integrated multi-omic approach to elucidate how specific rice genotypes mitigate CH_4_ emissions. The research highlights how the intersecting activities of H_2_-metabolizing, CH_4_-producing and other microbes in rhizosphere communities modulate CH_4_ cycling (Fig. 8).

**Fig. 8.**
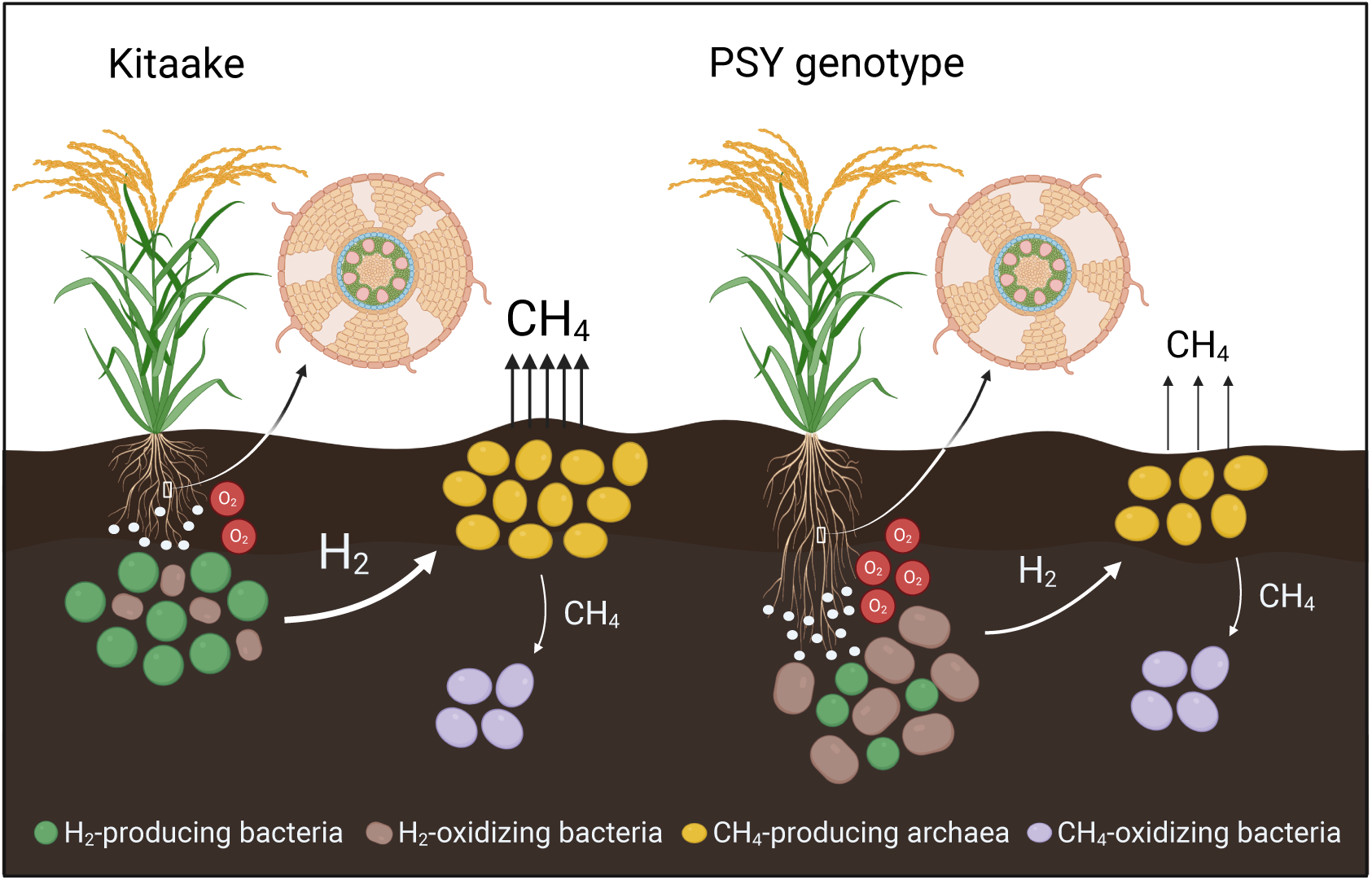
Proposed mechanism for the role of PSY rice genotypes mitigating CH_4_ emissions. PSY rice genotypes have longer roots with more aerenchyma compared with Kitaake. O_2_ is transported from the atmosphere to the rhizosphere through aerenchyma. PSY rice roots secrete more gluconeogenic acids (white circles), mostly amino acids, that can facilitate the activities of H_2_-oxidizing bacteria, thus reducing the total H_2_ pool for H_2_-dependent methanogenesis. The produced CH_4_ was partially oxidized *in situ* by methanotrophic bacteria and the other was emitted into the atmosphere. The figure is created with BioRender.com.

The CH_4_ production in our study was derived mainly from hydrogenotrophic methanogens (≥90%; Fig. 4c), consistent with extensive prior research^11,45,46^. Previous rice field studies using isotope-labeled substrates revealed that the presence of electron acceptors such as sulfate and nitrate channeled acetate towards CO_2_, and that CH_4_ was mainly produced from H_2_/CO_2_-dependent methanogenesis^47,48^. Dependence on H_2_ is likely explained by an insufficient supply of acetate. Acetoclastic methanogens (e.g., *Methanosarcina*) require 0.2-

1.2 mM acetate to activate CH_4_ formation^49,50^ whereas hydrogenotrophic methanogens (e.g., *Methanobacterium*) only need 2.8-10 Pa H_2_^51,52^. Low acetate concentrations may be due to limited acetate production or the inability of methanogens to compete with other organisms for acetate^47^. The transcriptional activities of nitrate and sulfate reductases in our study provide support for this hypothesis (Supplementary Figs. 11 and 12). These constituents (i.e., ∼2.0 mM of both nitrate and sulfate) were periodically added to water used to fertilize the rice plants during growth (Supplementary Table 2).

Although for all wild-type and transgenic rice genotypes the dominant pathway for CH_4_ production is H_2_-based methanogenesis, PSY1 and PSY2 genotypes, in two separate experiments, mitigated the cumulative CH_4_ emissions by up to 58%, relative to the Kitaake control (Fig. 2n). Such a reduction is higher than that achieved using a loss-of-function *gs3* rice allele reported in a recent study^11^. The most straightforward distinction between the wild-type and transgenic genotypes is the lower expression of *mcrA* and the lower ratio of transcripts for CH_4_ production versus oxidation (*mcrA*/*pmoB*) in microbiomes associated with the PSY1 genotypes (Fig. 4c). The much lower expression of *Methanocella mcrA* in the PSY1 genotypes compared to the Kitaake control (Fig. 4c) in the last growth stage is in line with the lower measured CH_4_ emissions (Fig. 2l). This difference may be partially attributed to higher O_2_ concentrations in the PSY rhizosphere due to more aerenchyma in the PSY compared with Kitaake roots (Fig. 1 j-l). Consistent with this, the activity of terminal oxidases in microbes associated with the PSY1 genotypes was higher than for microbes associated with the Kitaake control (Supplementary Fig. 3). Although *Methanocella* cells encode genetic machinery to resist oxidative stress and can persist in aerated environments^53^, their methanogenesis activity would be inhibited by oxygen exposure until conditions become anoxic^54^.

Another factor that may have affected CH_4_ production is the availability of the methanogenesis substrate, H_2_. In rice paddies, H_2_ is typically produced during microbial fermentation, e.g., of plant-derived sugars, and can be consumed via diverse processes^55^. We found much higher activities of H_2_-consuming hydrogenases but lower activities of H_2_-producing counterparts associated with the PSY1 genotypes (Fig. 5). This likely reduced the H_2_ available for hydrogenotrophic methanogenesis, compared to that with the Kitaake.

Activities of the root-associated H_2_-metabolizing microorganisms can be altered by rice root characteristics. As larger root systems can be associated with increased CH_4_ emissions in paddy fields^56^, rice breeding efforts for mitigating CH_4_ emissions have aimed to generate plants with smaller root systems^10,11^. However, overexpression of rice *PSY* genes led to longer roots (Fig. 2k) and only the PSY1 genotype displayed reduced root biomass (Fig. 2i), compared with the control. The distinct developmental phenotypes observed in rice plants overexpressing *OsPSY1* and *OsPSY2* may be driven by differences in receptor binding and tissue-specific expression. Similar behavior has been observed in other small peptide families, such as the CLE family^57^. Still, both OsPSY1 and OsPSY2 genotypes appear to share common molecular mechanisms contributing to comparable reductions in CH₄ emissions (Fig. 2k-n).

Given the similar reduction in CH_4_ emissions despite differences in PSY1 and PSY2 root biomass, we considered whether the enrichment of PSY exudates in gluconeogenic acid metabolites (mostly amino acids) could alter rhizosphere microbial H_2_ metabolism, with implications for CH_4_ production (Supplementary Table 6). H_2_-consuming microorganisms that are more invested in use of these compounds than H_2_-producing bacteria had significantly higher activities associated with the PSY1 genotypes (Fig. 6). Further, metabolic modeling results indicate that PSY1 over-secreted compounds suppress H_2_ production activity (Fig. 7).

Intriguingly, the modeling predicts the increase in biomass of H_2_-producing bacteria despite the decreased H_2_ generation (Supplementary Fig. 10). This may be explained by the high relative abundance of amino acids in PSY1 exudates (Supplementary Table 6). Root-associated H_2_-producing bacteria may partly rely on extracellular amino acids that can be converted to pyruvate and acetyl-CoA, and thus rely less on fermentation of sugars to form H_2_^58,59^. The rice-derived amino acids could also alleviate the necessity for microbial nitrogen fixation (Supplementary Fig. 4), reducing the formation of byproduct H_2_^60^. Conversely, microbial H_2_ consumption was barely affected by the PSY1 over-secreted compounds (Fig. 7). Overall, the distinct exudate profiles of PSY1 rice genotypes inhibit the activities of H_2_-producing microorganisms but stimulate that of the H_2_-consuming counterparts, drawing down the pool of H_2_ available for methanogenesis thus mitigating final CH_4_ emissions (Fig. 8).

It has been demonstrated that exudation rate and composition vary throughout plant development^61^. Extending the exudation studies to later developmental stages of rice plants will be informative as the current analysis includes relatively early developmental stages. Such future research will further advance our understanding of the molecular mechanisms through which PSY1 affects plant-microbe interactions and associated biogeochemical cycles.

## Methods

### Plant materials and growth conditions

*Oryza sativa* ssp. *Japonica* cultivar Kitaake was used throughout this study as the wild-type (wt) background. Rice seeds were dehulled, surface-sterilized in 20% bleach for 30 minutes, and thoroughly washed with autoclaved water. The seeds were grown in 1x Murashige and Skoog (1xMS) salt mixture with 1% sucrose (pH 5.65), either as a solid medium containing 0.3% Gellex (Gellan Gum, Caisson Laboratories) or hydroponically in flasks with 100 mL of the same media without Gellex. For solid medium conditions, seeds were sown on plates, sealed with Micropore tape for gas exchange, and placed vertically for 7 days. For hydroponic conditions, seeds were first germinated on plates with water for 2 days in darkness, then transferred to flasks for another 5 days. The incubators used for rice germination and growth were set with a 14-h-light/10-h-dark photoperiod at 28°C/24°C.

For synthetic peptide treatments in hydroponic conditions, peptide (or water for untreated plants) was added to the MS media before transferring the plants. The length and concentration of the peptide treatment are specified in the figure legends. The synthetic OsPSY1 peptide was obtained from Pacific Immunology (Ramona, CA, USA), is tyrosine sulfated, and was diluted in ddH2O to a final concentration of 1 mM. For each experiment, MS media was freshly prepared and cooled for 1 hour in a 55-60°C water bath after autoclaving before adding chemicals.

For primary root elongation, seedlings grown hydroponically or in solid media were photographed 7 days after germination, and the primary root length was measured with Fiji Is Just ImageJ^62^.

### Plasmid construction and rice transformation

DNA constructs were created with the Gateway cloning technology^63^. The genomic OsPSY1 and OsPSY2 sequences were amplified using the primers described in Supplementary Table 9. These sequences were then recombined with pENTRTM/D-TOPO (Invitrogen, Cat#45-0218) to yield pTOPO_OsPSY1 and pTOPO_OsPSY2. The latter vectors were used in a Gateway LR cloning (Gateway® LR ClonaseTM II Plus Enzyme Mix, Invitrogen; Cat#:12538-120) with pC1300-Ubi-Nos to yield *Ubi::OsPSY1* and *Ubi::OsPSY2* constructs. The generated vectors were transferred to *Agrobacterium tumefaciens* strain EHA105 and then introduced into the rice lines via Agrobacterium-mediated transformation. Rice transformation was performed following previously published protocols^64^. Post-transformant screening was based on hygromycin resistance and PCR determination of T-DNA insertion. Out of 52 positive lines for *Ubi::OsPSY1* and 53 for *Ubi::OsPSY2*, four homozygous lines were selected for characterization. Eventually, three homozygous lines, *Ubi::OsPSY1#9* (PSY1-9), *Ubi::OsPSY1#17*(PSY1-17), and *Ubi::OsPSY2#34* (PSY2-34), were selected for detailed studies.

### Gene expression analysis by quantitative PCR (qPCR)

Total RNA (1mg) was extracted from the roots of seven-day-old rice seedlings grown hydroponically using TRIzol reagent (Invitrogen) and treated with the TURBO DNA-free kit (Ambion) to remove residual genomic DNA. cDNA was synthesized using the High-Capacity cDNA Reverse Transcription Kit (Applied Biosystems). The cycle threshold (Ct) value was measured on a Bio-Rad CFX96 Real-Time System coupled to a C1000 Thermal Cycler (Bio-Rad) using the iTaq Universal SYBR Green Supermix (Bio-Rad). Normalized relative quantities (NRQs) were obtained using the qBase method^65^, with LOC_Os08g18110 (putative alpha-soluble NSF attachment protein) and LOC_Os11g26910 (putative SKP1-like protein 1B) as reference genes for normalization across samples^66^. NRQ values were normalized to the mean value obtained in wild-type plants. NRQ Melting curve analyses at the end of the process and “no template controls” were performed to ensure product-specific amplification without primer–dimer artifacts. Primer sequences are given in Supplementary Table 9. Three biological replicates were analyzed.

### Cellular analysis using LSCM

Laser scanning confocal microscopy (LSCM) was performed throughout the study using a Plan Apochromat 63x/1.40 oil and 20X/0.75 CS2 lens on a Leica TCS SP8 microscope. Cell wall staining, seedling fixation and staining were performed using an adapted Clearsee protocol^67^. Briefly, the primary root of 7-day-old seedlings was sectioned into three short segments with a length of 0.5cm starting from the root tip. The root segments were fixed for 1h at room temperature in 10% neutral buffered formalin in PBS, using 6-well plates, then washed five times for 1min with PBS 1X. For root transverse section analysis, the 1 cm root fragments collected 4cm from the root tip were embedded in 4% agarose and sectioned using a Leica automated vibrating blade microtome (VT1200 S). Once fixed, root fragments and transverse sections from those fragments were cleared in Clearsee solution for at least 24 hours under mild shaking. Fixed and cleared samples were incubated overnight in a Clearsee solution supplemented with 0.1% Calcofluor White. After 12 hours, the staining solution was removed, and samples were rinsed once in fresh Clearsee solution, then washed twice for at least 120 minutes in a renewed Clearsee solution with gentle shaking. Roots were carefully placed on a microscope slide with ClearSee and covered with a coverslip. Excitation and detection windows were set as follows: Calcofluor white for excitation at 405 nm, and detection between 415 and 570 nm. Central longitudinal section images were acquired from root complete fragments. Mature cortical cell length was assessed in 10 consecutive cells in the second cortex layer. Transverse section images were acquired to quantify percentage aerenchyma formation.

### Soil collection and growth conditions in the greenhouse experiments

All data presented in this article were generated from two greenhouse studies carried out at the University of California—Davis in the fall of 2022 (EXP1) and the spring-summer of 2024 (EXP2). Soil from a rice field in Davis (38.543224 degrees North and -121.810536 West) was collected in July 2022 (for EXP1) and in July and December 2023 (for EXP2) using a front-end loader to gather down to a depth of approximately 6-10 inches. Soil samples were analyzed at the UC Davis Analytical Lab for chemical content (Supplementary Table 10). For both experiments, all soils were transported back to the greenhouse, crushed to a 2mm particle size, thoroughly homogenized, and stored until planting. Then, the soil was scooped into new 5.5 x 5.5 inch pots (2000g per pot), which were then placed into 5 23 gallon tubs per genotype (18 pots each). Prior to seedling transplantation, enough water was added to the tubs to submerge the soils.

Seeds were germinated in solid MS media, and root growth was followed as previously described for 7 days. Axenic seedlings were then transplanted into the watered pots (one seedling per pot) in the greenhouse, where they were irrigated with deionized water every other day to keep the soil submerged. Fertilized water [N, 77 ppm (parts per million); P, 20 ppm; K, 75 ppm; Ca, 27 ppm; Mg, 17 ppm; S, 68 ppm; Fe, 1.5 ppm; Cu, 0.02 ppm; Mn, 0.5 ppm; Mo, 0.01 ppm; Zn, 0.1 ppm; B, 0.5 ppm] replaced the distilled water in the tubs after microbiome samples were collected on day 15, 30, 55 and 70 after transferring to the pots. Each tub contained only one plant genotype to avoid root exudate compositional changes. We also kept a separate tub with unplanted pots or bulk soil control. Tubs were distributed in the greenhouse benches following a completely randomized block design. All weeds were manually removed from the pots when identified.

In experiments 1 and 2, plant developmental stage and height were measured on the microbiome samples collection days.

In addition, in experiment 2, roots and shoots were collected 100 days after transferring plants to pots, and dry biomass was measured. Seed number and one thousand seed weight were also calculated for the same plants.

### Cell wall residue preparation

Samples for lignin analysis were collected from 7-d-old rice seedlings or from root systems of 90-d-old plants grown in agricultural soil. Samples from older root systems were collected in the same root-shoot junction location as the one used for rhizosphere sample collection.

The fresh root biomass was lyophilized for three days and then ball milled biomass was extracted sequentially by sonication (15 min) with 80% ethanol (three times), acetone (one time), chloroform–methanol (1:1, v/v, one time) and acetone (one time).

### TGA lignin

Aliquots of 15 mg of cell wall residue were weighed into Eppendorf tubes and mixed with 1 ml of 3 N HCl and 0.1 ml thioglycolic acid (adapted from the reference^68^). Samples were incubated at 80°C for 4 hours and repeatedly mixed. Samples were rapidly cooled on ice and centrifuged for 10 min at 15,000g. The supernatant was discarded. Pellets were washed 3-times with distilled water (1 ml). Thereafter, pellets were incubated with 1 ml of 1 N NaOH for 16 hours on a shaker at room temperature. The suspension was centrifuged for 10 min at 15,000g. The supernatant was carefully transferred into a 2-ml Eppendorf tube. The resulting supernatant was combined with the first alkaline supernatant and mixed with 0.2 ml concentrated HCl. Samples were incubated for 4 hr at 4°C to precipitate the lignothioglycolate derivates. The samples were centrifuged, the supernatant discarded, and the pellet solubilized in 1 ml of 1 N NaOH. Absorbance of the resulting solution was measured at 280 nm. Results were expressed as an absorbance per mg cell wall.

### Measurement of methane emissions

Gas flux measurements of CH_4_ were made in real-time in a transparent dynamic flow chamber using Picarro G2508 Gas Analyzer. In the fall 2022 experiment, a total of 4 timepoints (15d, 30d, 55d, 70d) were taken across the growing season for 5 biological replicates corresponding to Kitaake, PSY1-17, PSY1-9, and bulk soil sample types. In the spring-summer 2024 experiment, a total of 12-time points were taken across the growing season for 5 biological replicates corresponding to Kitaake, PSY1-17, PSY2-34, and bulk soil sample types. Each replicate constituted a single rice plant except for bulk soil samples, which represented a negative control. At the start of every measurement, a chamber was placed onto a rice plant with minimal disturbance. The chamber was left on for at least 5 minutes for flux analysis. Gas fluxes were calculated by fitting linear regression lines across measurement time windows.

The conversion to mass fluxes using Ideal Gas Law equation:

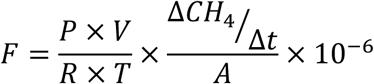

where F is the CH_4_ flux (mg m^-2^ h^-1^), P is the pressure, V is the volume, and A is the area of the sampling chamber. R is the ideal gas constant, and △CH_4_/△t is the slope of the gas concentration change over timefitted by a linear equation. T is the temperature in the sampling chamber (K).

### Microbiome sample collection

Rhizosphere samples were collected 15, 30, 55, and 70 days after rice was transplanted into pots, following the protocol described before^35^. Briefly, we pulled out rice roots from pots and manually shook them to remove excess soil from the roots, leaving only ∼1 mm of soil attached on roots. Then we put roots into 50-mL Falcon tubes that contained 15 mL of Powerbead Solution (Qiagen), stirred vigorously with sterile forceps to obtain closely attached soil, and frozen immediately in liquid nitrogen after removing roots. Samples were stored in the - 80°C freezer before nucleic acid extraction.

### Extraction and sequencing of total DNA and RNA

Approximately 2.0 g of rhizosphere samples were used for co-extraction of DNA and RNA by combining the Qiagen RNeasy PowerSoil Total RNA Kit and RNeasy PowerSoil DNA Elution Kit. Isolated nucleic acids were qualified and quantified using Nanodrop (Thermo Fisher) and Qubit (Invitrogen). For DNA samples, library was prepared using Roche KAPA HyperPrep Kit and sequenced on Illumina NovaSeq6000 PE150 platform in Maryland Genomics. For RNA samples, ribosomal RNA (rRNA) was first depleted using the NEBNext rRNA Depletion Kit, then library was prepared using NEBNext Ultra II Directional RNA Library Prep Kit and sequenced on the same above platform. In total, 72 DNA samples and 72 RNA samples including replicates were sequenced, generating 974.8 Gbp and 920.7 Gbp of metagenomic and metatranscriptomic raw read data. Details for each sample can be found in Supplementary Table 1.

### Metaomics *de novo* assembly and binning

Raw reads from metagenomic sequencing were processed using BBDuk contained in the BBTools suite (https://jgi.doe.gov/data-and-tools/software-tools/bbtools/). Biological replicates from the same samples were merged using BBMerge. Samples were individually assembled using metaSPAdes (v3.15.5) with default parameters^69^. Scaffolds longer than 1 kb were binned by a combination of MetaBAT2 (v2.15)^70^, CONCOCT (v1.1.0)^71^, MaxBin2 (v2.2.7)^72^, and VAMB (v3.0.2)^73^, using differential abundance calculated by pairwise cross-mapping against all samples by BBMap. Resulting bins from the same samples were integrated and optimized using DAS Tool (v1.1.2)^74^. Optimized bins derived from all samples were dereplicated at 98% whole-genome average nucleotide identity (ANI) using dRep (v3.4.0)^75^, generating a set of approximately subspecies-level genomes. Quality of representative genomes was assessed using CheckM2 (v1.0.1)^76^ and taxonomy was assigned by GTDB-Tk (v2.3.0)^77^ based on the reference database version r214^78^ and further converted to NCBI taxon. Only genomes with predicted completeness ≥50% and contamination ≤5% were used for downstream analyses (n=296).

For metatranscriptomics, raw reads were pre-treated using BBDuk, and rRNA was filtered by SortMeRNA (v4.3.6)^79^. Clean reads primarily containing mRNA were individually assembled using rnaSPAdes (v3.15.5)^80^ to generate transcripts.

### Gene prediction and annotation

Open reading frames (ORFs) in assembled scaffolds and transcripts were predicted by Prodigal (v2.6.3)^81^. Protein sequences from all samples were combined together and dereplicated at 95% ident along with 85% coverage to generate a non-redundant gene set, using Linclust integrated in MMseqs2^82,83^. Predicted genes were basically annotated against KEGG^84^, UniRef100, and UniProt^85^ databases by USEARCH v10^86^, and against the CAZy database^87^ using the online server following the manual instruction^88^. Protein domains were searched against Pfam (release 36.0)^89^ using Astra (https://github.com/jwestrob/astra).

For the identification and clustering of *rpS3* genes, the PF00189 HMM was first used to search against predicted proteins based on pre-built noise cutoffs using HMMER (v3.3)^90^. Corresponding nucleotide sequences of protein hits were pulled out and clustered at 99% ident^91^ using VSEARCH (v2.13.3)^92^. Representative sequences from each cluster were blasted against Genbank nr database^93^ to get preliminary taxonomic information. Precise taxonomy was further retrieved by phylogeny construction that involved alignment of sequences using MAFFT (v7.453)^94^, trimming of alignments using trimAl (v1.4.rev15)^95^, and generation of phylogenetic trees using IQ-TREE v1.6.12 with best-fit models automatically determined^96^. Trees were decorated on the iTOL server for better visualization^97^.

For genes closely related to CH_4_ cycling, the PF02249 and PF02745 HMMs were used to recruit McrA-like sequences; the TIGR03079 was to PmoB-like sequences; and the PF02964 was to the gamma subunit of methane monooxygenase hydroxylase. Further identification and classification were performed based on phylogeny as described above.

For genes involved in H_2_ production and consumption, three curated HMMs were used to target sequences homologous to [NiFe], [FeFe], and [Fe] hydrogenases, respectively^38^. Groups and subtypes of identified hydrogenases were distinguished based on protein phylogeny. [FeFe] Group A1 hydrogenases were further classified using the online tool HydDB which integrates genetic context for accurate classification^38^. For genes related to nitrate reduction, proteins that contain molybdopterin-binding domain or molybdopterin oxidoreductase domain were phylogenetically classified into distinct groups, including NarG and NapA exclusive groups. NifH proteins were identified using the TIGR01287 HMM that targets nitrogenase iron protein included in all three nitrogenase forms (i.e., Mo-Fe, V-Fe, and Fe-only). Reductive DsrA was identified by DiSCo (v1.0.0)^98^. Taxonomy of genes was identical to that of gene-located scaffolds which was assigned if 50% of the coexisting proteins belonged to the same taxon.

### Abundances of assembled genomes and genes

Metagenomic abundances of genomes and genes were calculated using CoverM (v0.6.1, https://github.com/wwood/CoverM). Given genomes dereplicated at 98% ANI, a minimum read identity during mapping was set at 98% together with a minimum aligned percent of 75% in the “genome” mode. For non-redundant genes at 95% ident level, corresponding scaffolds were mapped by reads with a minimum identity of 95% and aligned percent of 75% using the “contig” mode. Abundances were considered to be zero if the covered fraction by any read was <50%. Normalized abundances were calculated by dividing coverage by total read length and multiplying a factor of 1E11 for visualization.

Overall metatranscriptomic abundances of genomes were calculated using CoverM with the same cutoff parameters as above, followed by a normalization in which aligned read counts for each genome were divided by total read size and genome length, and multiplied by a factor of 1E15. To calculate the abundances of genes, corresponding scaffolds were first mapped by metatranscriptomic reads using Bowtie2 with “--end-to-end” “--sensitive” mode^99^. Exact aligned reads for genes inside scaffolds were counted using featureCounts (v2.0.6)^100^. Final abundances, that said transcriptional activities, were normalized by dividing read counts by gene length and multiplying a factor of 1E11 for visualization.

### Enrichment of genes/pathways in up-regulated genomes

Based on calculated overall transcriptional abundances of genomes, DESeq2^101^ was used to identify significant differences in genome activities between the Kitaake and PSY1 genotypes, and between rice planted and unplanted groups. We consider genomes as “up-regulated genomes” if: 1) they had significantly higher overall activities (log2FoldChange ≥ 1, p ≤ 0.05) in PSY1 rice genotypes compared with the Kitaake in ≥ 1 rice growth stage; 2) they never showed significantly higher overall activities in the Kitaake than PSY1 genotypes; and 3) they were more active in rice planted groups than in bulk soils. Finally, 17 of 296 genomes were identified as up-regulated genomes in PSY1 genotypes.

KEGG Orthology (KO) identifiers were assigned to predicted proteins in all genomes, using kofamscan (v1.3.0; https://github.com/takaram/kofam_scan) with KOfam set r2023-10-02. Only hits above score thresholds and with the highest score were selected for individual proteins. Next, we performed Fisher’s exact test to evaluate the over/under enrichment of identified KOs in up-regulated genomes (n = 17) relative to remaining genomes (n = 279). A total of 3509 KOs that appeared in ≥ 10 genomes were tested, and the resulting p values were adjusted using Benjamini-Hochberg method^102^. KOs with FDR ≤ 0.05 were considered significantly enriched (odds ratio > 1) or depleted (odds ratio < 1) in up-regulated genomes.

Given KEGG metabolic pathways comprised of multiple KOs, we quantified pathways by summing up abundances of all genes that are associated with related KOs. Differential abundances were calculated by subtracting abundances in the PSY1 genotypes from abundances in the Kitaake, shown as PSY1-17 and PSY1-9 respectively.

### Calculations of genome-wide SAP values

We followed the previous paper to predict SAP values from genomes^42^. Briefly, we first calculated the relative abundances (number of genes normalized by the total number of genes in genomes) of sugar and acid genes that are implicated in the corresponding degradation pathways, based on the list in the above reference. The generalized linear model was then used to predict SAP values for each genome, as:

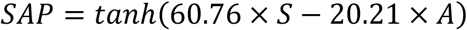

where S and A indicate relative abundances of sugar and acid genes, respectively. Statistical difference in average SAP values between genome groups was calculated using the Wilcoxon rank sum test.

### Rice roots and exudate collection

Samples for metabolomic analysis were collected from 7-day-old rice seedlings grown vertically on 1x MS solid media plates. Prior to collection, seedlings were removed from the plates and thoroughly washed with deionized water to remove any residual media from the roots. For root tissue samples, the entire root system of 20 seedlings was ground using a mortar and pestle, with a subsample transferred to a 2 mL tube (Eppendorf #022363352). Tissue fresh weight was recorded with pre-weighted tubes, and samples were stored at −80 °C until further processing. For root exudate samples, 15 seedlings were placed in 50 mL flasks containing ultra-pure water and held in place by sponges so that only the root tips were submerged. The flasks were incubated with shaking for 2 hours before the exudates were collected in 50 mL tubes (Fisher Scientific#14-959-49A). Exudates were initially stored at −20°C for a few hours, then transferred to −80°C until further use.

### Liquid chromatography sample preparation

To extract metabolites from 300-400 mg root tissue and agar control media, samples were first frozen, lyophilized dry (FreeZone 2.5 Plus; Labconco), then powderized by bead-beating (MiniBeadbeater96, BioSpec) for 5 seconds, 2x, using a 3.2 mm stainless steel bead. Next, 500 uL of 100% methanol was added to each powderized root sample and 1.5 mL to each powderized agar sample, briefly vortexed, then sonicated in an iced water bath for 10 minutes. Samples were centrifuged (5 min, 10k rpm) to pellet debris, supernatant removed, and transferred to a 2 mL Eppendorf, dried in a SpeedVac (SPD111V; Thermo Fisher Scientific). To extract metabolites from exudates collected in 40 mL water, samples were first frozen then lyophilized dry. Next, 3 mL of 100% methanol was added to each sample, briefly vortexed, then sonicated in an iced water bath for 10 minutes. Samples were centrifuged (5 min, 10k rpm) to pellet debri, supernatant removed and transferred to a 5 mL Eppendorf, then dried in a SpeedVac. Next, 1 mL of 100% methanol was added to each dried extract, briefly vortexed, then sonicated in an iced water bath for 10 minutes. Samples were centrifuged again, then supernatant removed and transferred to a 2 mL Eppendorf and dried in a SpeedVac. All dried extracts were stored frozen at - 80C. The same extraction procedures were followed using empty 50 mL and 2 mL tubes to serve as extraction controls, and a 50 mL tube containing 50 mL water to serve as an exudate collection control (3 replicates each).

In preparation for LC-MS, samples were resuspended in 100% methanol containing a mix of isotopically labeled internal standards (dx.doi.org/10.17504/protocols.io.kxygxydwkl8j/v1). Resuspension volume was varied for each root extract to normalize by amount of extracted biomass (1500 mg/mL), while all other samples were resuspended in 150 uL.

### LC-MS/MS metabolomics analysis

Samples were run using both normal and reverse phase chromatography. Each of these was performed using an Agilent 1290 LC stack, with MS and MS/MS data collected using an Exploris 120 Orbitrap MS (Thermo Fisher Scientific, San Jose, CA) for normal phase, and QExactive HF Orbitrap for reverse phase. Mass spectrometer parameters and gradients are provided in this protocols.io (dx.doi.org/10.17504/protocols.io.kxygxydwkl8j/v1) for normal phase, and this protocols.io for reverse phase (ref here). For both normal phase and reverse phase, full MS1 spectra was collected at 60k resolution in both positive and negative ionization mode, with MS/MS fragmentation data acquired using stepped then averaged 10, 20, and 40 eV collision energies at 15k resolution. For normal phase, the MS1 m/z range for data collection was 70-1050, while for reverse phase, both m/z 80-1200 and m/z 1000-3000 were used in separate runs to cover mass ranges for metabolites and peptides. Samples consisted of three biological replicates each and extraction controls, with sample injection order randomized by replicate and an injection blank of 100% methanol run between each sample, with the blank replaced by an injection of internal standard mix every third sample, as well as QC mix every 15 samples. The raw data from the metabolite LC-MS runs are provided in Supplementary Dataset 1. For untargeted data analysis, features with 0.8 min<RT<18.5 min (HILIC column) and significant (adj p-value < 0.05) maximum peak height fold-change between sample and extraction control >3 were selected. We performed a Feature-Based Molecular Networking (FBMN)^103^ workflow using MZmine-3.7^104^ and a Global Natural Products Social Molecular Networking (GNPS2; http://gnps2.org)^105^. The MZmine workflow was used to generate a list of features obtained from extracted ion chromatograms containing chromatographic peaks within a narrow m/z range and filtered to remove isotopes. For each feature, the most intense fragmentation spectrum was uploaded to GNPS for putative identification by comparison with mass spectra deposited in the database. Compound classes were attributed to identified compounds using NPClassifier^106^.

For targeted analysis, compound identification was performed by comparing detected *m/z*, retention time (RT) and fragmentation pattern to standards run in-house using the same LC-MS methods and assigning a confidence level based on these criteria. (Supplementary Dataset 3; Supplementary Table 5). For each identified compound, the highest level of confidence (Exceeds Level 1)^107^ was achieved when the measured m/z agreed with the expected theoretical m/z with less than 5 ppm error, detected RT of the feature was within 0.5 minutes of theoretical, and fragmentation pattern matched that of the standard. When MSMS was not collected, some compounds were identified based on only mz and RT (Level 1); mis-matching MSMS would invalidate an identification.

### Construction and analysis of genome-scale metabolic models

Microbial genomes that have hydrogenase genes and predicted completeness >90% and contamination <5% were annotated using the App “Annotate Genome/Assembly with RASTtk” (v1.073) in KBase^108^. The resulting annotated genomes were used as inputs for the App “MS2 - Build Prokaryotic Metabolic Models with OMEGGA” (v.2.0.0) to build gap-filled metabolic models using media comprised of basic nutrients and PSY1 over-secreted compounds (Supplementary Table 8). Basic nutrients included glucose and glutamate as primary carbon sources, which were present and had no significant difference in concentrations across rice genotypes. Also included were standard essential factors for microbial growth (e.g., Fe^2+^). PSY1 over-secreted compounds were measured at significantly higher concentrations in root exudates from PSY1 rice genotypes compared with the Kitaake.

Dynamic flux balance analysis (dFBA) was performed for each microbial model individually, assuming well-mixed batch growth conditions, using the Computation Of Microbial Ecosystems in Time and Space (COMETS) platform (v2.12.2)^109^. We set the initial biomass at 1 × 10^−6^ g and used default simulation parameters. To compare results across varying conditions, we calculated the maximal potential of substrate turnover and biomass yield when microbial growth stopped and substrates no longer changed, for each condition. As glucose and glutamate in basic nutrients were likely the most abundant carbohydrate and amino acid secreted by rice roots^110,111^, concentrations of PSY1 over-secreted compounds were varied from 0.01-1 folds as that of basic nutrients. O_2_ is transported to the rhizosphere through rice aerenchyma and its concentrations were set at 1-10 folds as that of basic nutrients. H_2_ concentrations were set at 0 when testing H_2_ production potential and at 10 folds when testing H_2_ consumption potential. For visualization purposes, we calculated the consumed H_2_ amounts by subtracting the final values from the initial.

### Statistical analysis

NMDS analyses for the rpS3 gene were performed and plotted using “vegan”^112^ and “ggplot2”^113^ packages in R (v4.2.2)^114^. Differences and significance between groups were assessed by analysis of similarities (ANOSIM) and permutational multivariate analysis of variance (PERMANOVA) tests. Outliers were detected and removed from replicates using the median absolute deviation (MAD) test^115^. Differential expressions of genomes between samples were calculated by DESeq2^101^.

## Data availability

All metagenomic and metatranscriptomic sequencing reads have been deposited in NCBI Sequence Read Archive (SRA) with the project accession number: PRJNA1161606. Prior to publication, assembled microbial genomes can be accessed at: https://ggkbase.berkeley.edu/Assembled_genomes_from_rice_genotype_experi ment. Please note that it is necessary to sign up as a user (simply provide an email address) in order to view and download the data. The submission of metabolomics raw data is in progress.

## Acknowledgments

We would like to thank Ryan C. Packer and Rory M. Greenhalgh for their help setting up the first greenhouse experiment. We thank Bethany C. Kolody and Jackie Zorz for the assistance in using R, and Daniel A. Gittins for the help in hydrogenase HMMs. We thank Jennifer Pett Ridge for the helpful suggestions.

Funding for this research was provided by the Chan Zuckerberg Initiative. This material is based upon work supported by the Joint BioEnergy Institute, U.S. Department of Energy, Office of Science, Biological and Environmental Research Program under Award Number DE-AC02-05CH11231 with Lawrence Berkeley National Laboratory. Funding was also provided by the Bill and Melinda Gates Foundation (grant number INV-037174 to J.F.B.). The findings and conclusions are those of the authors and do not necessarily reflect positions or policies of the Bill and Melinda Gates Foundation. The metabolomic analysis was conducted by the U.S. Department of Energy Joint Genome Institute (https://ror.org/04xm1d337), a DOE Office of Science User Facility supported by the Office of Science of the U.S. Department of Energy operated under Contract No. DE-AC02-05CH11231. A.M.S. is supported by a U.S. Department of Agriculture (USDA) National Institute for Food and Agriculture (NIFA) Postdoctoral Fellowship (2023-67012-39889).

## Author contributions

The study was conceived by L.D.S., M.F.E., P.C.R., and J.F.B. M.F.E., S.S, A.T.A.J., T.S.W., and A.M.S. processed the soil from rice paddies at UC Davis, planted rice in the greenhouse, and performed phenotypic analyses for EXP1 and EXP2. L.D.S. and J.K. measured CH_4_ flux and collected rhizosphere soil samples with the help of M.F.E., A.T.A.J., and A.M.S in EXP1. M.F.E., S.S., A.T.A.J., and T.S.W measured CH_4_ flux in EXP2. L.D.S. extracted DNA and RNA from collected soil samples and sequenced them and performed all metagenomic and metatranscriptomic analyses. M.F.E. and T.S.W. collected rice root tissue and exudate samples for metabolomic analysis. T.R.N., K.B.L. and B.P.B. designed the metabolomics study, generated and processed the metabolomic data, and M.F.E. analyzed the metabolomic data. H.T. performed the lignin analysis in young and adult rice plants. H.V.S. supervised cell wall analysis. L.D.S. constructed microbial metabolic models and performed dynamic flux balance analysis with the help of I.D. and D.S. R.S. provided technical support in processing sequencing data. L.D.S., M.F.E., P.C.R. and J.F.B. wrote the manuscript with input from all the authors.

---

**Supplementary Fig. 1.**
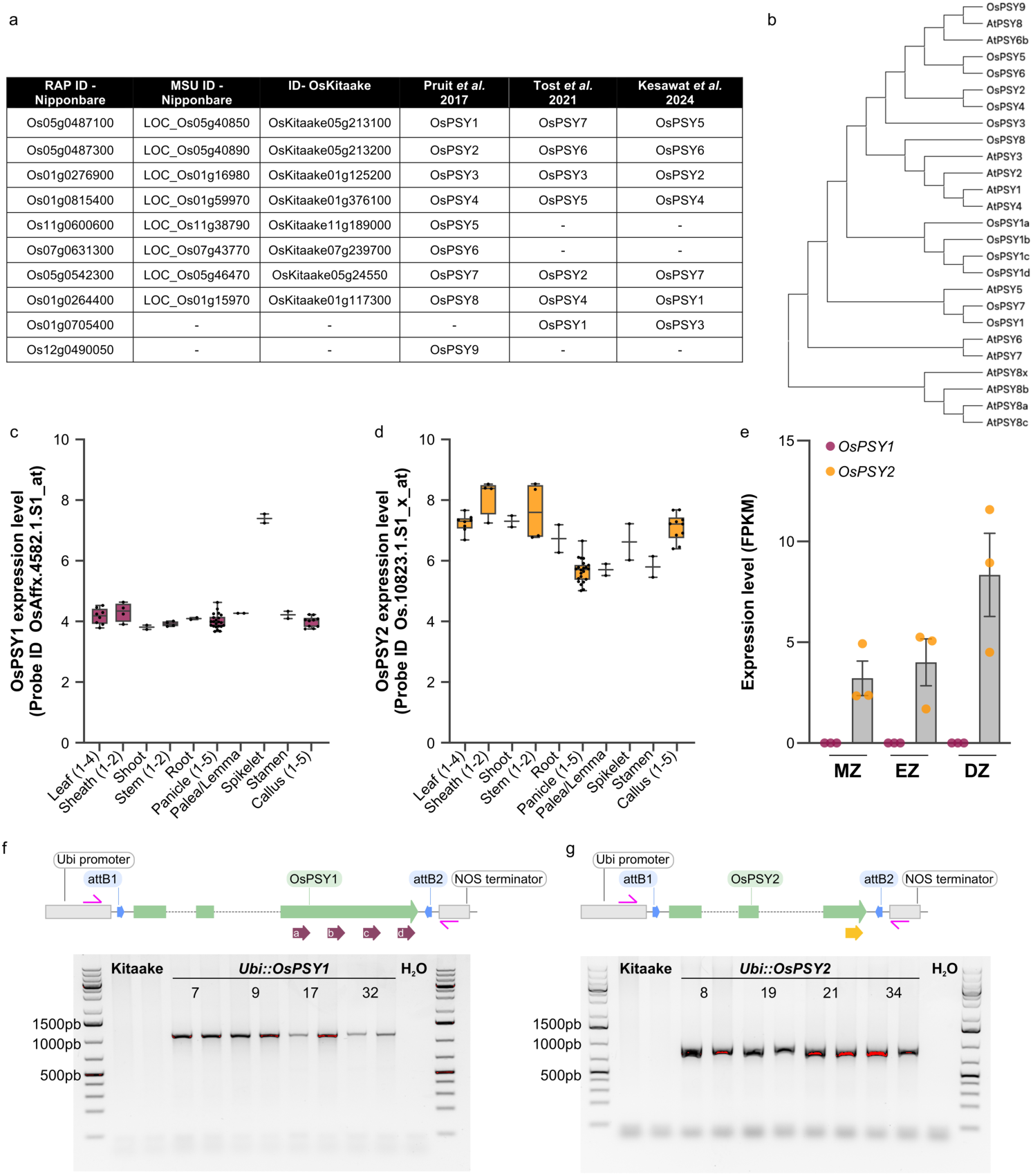
PSY peptide family in rice. (**a**) Table summarizing locus IDs and gene names assigned to different OsPSY genes in rice. In this manuscript, we followed the naming convention from Pruitt et. al, 2017. (**b**) Phylogenetic tree showing the relationships between PSY homologs in *Arabidopsis thaliana* (AtPSYs) and *Oryza sativa* (OsPSYs). Expression levels of *OsPSY1* (**c**) and *OsPSY2* (**d**) across mature organs in cultivar Minghui 63. Normalized microarray expression values were plotted using data from probe IDs OsAffx.4582.1.S1_at and Os.10823.1.S1_x_at, respectively, from NCBI SRA BioProject PRJNA120617. (**e**) Expression profiles of *OsPSY1* and *OsPSY2* along the root longitudinal axis of 4-d-old rice seedlings using data extracted from publicly available data sets. MZ = meristematic zone, EZ = elongation zone, and DZ = differentiation zone. (**f, g**) Diagrams illustrating the gene structure and cloning design of OsPSY1 (**f**) and OsPSY2 (**g**) in the pC1300 vector. The regions highlighted in pink mark the oligos for the Ubiquitin promoter and NOS terminator (Supplementary Table 1) used to validate construct insertion into the rice genome. Below the diagrams, agarose gels show PCR amplification products of 1156 bp in *Ubi::OsPSY1* (#7, #9, #17, and #32) plants and 897 bp in *Ubi::OsPSY2* (#8, #19, #21, and #34) plants. These bands are absent in the non-transgenic Kitaake control plants.

**Supplementary Fig. 2.**
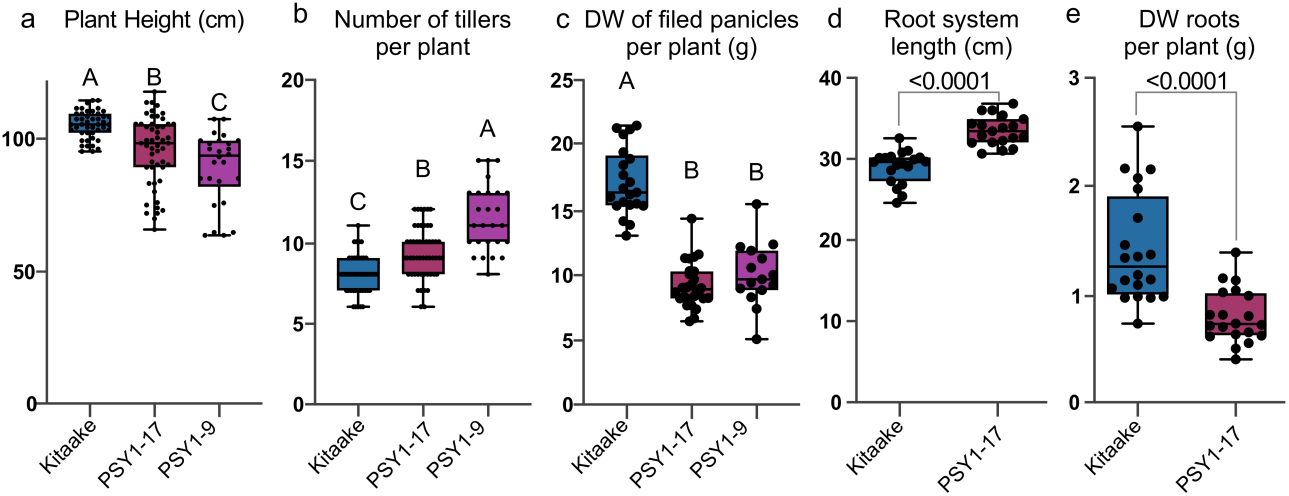
Phenotypic profiling of EXP1. Phenotypic profiling of Kitaake, PSY1-17, and PSY1-9 grown in EXP1 including (**a**) plant height (cm), (**b**) number of tillers per plant, (**c**) DW of filled panicles per plant. Phenotypic measurements were taken 70 days after transplanting for (**a**) (n=50 plants for Kitaake and PSY1-17, n=30 plants for PSY1-9),(**b**) (n=50 plants for Kitaake and PSY1-17, n=30 plants for PSY1-9), while (**c**), was analyzed at the end of the plant life cycle 100 days after transplanting (n=20 plants). (**d**) Root system length and (**e**) dry weight (DW) of the roots per plant from Kitaake and PSY1-17 of rice plants 30 days after transplanting in EXP1 (n=20 plants). The data shown in this figure are box and whisker plots combined with scatter plots; each dot indicates the measurement of the designated parameter listed on the y-axis of the plot. In (**a**), (**b**), and (**c**) different letters indicate significant differences, as determined by one-way ANOVA followed by Tukey’s multiple comparison test (P < 0.05). In (**d**) and (**e**), P values are calculated by two-tailed Student’s t-test.

**Supplementary Fig. 3.**
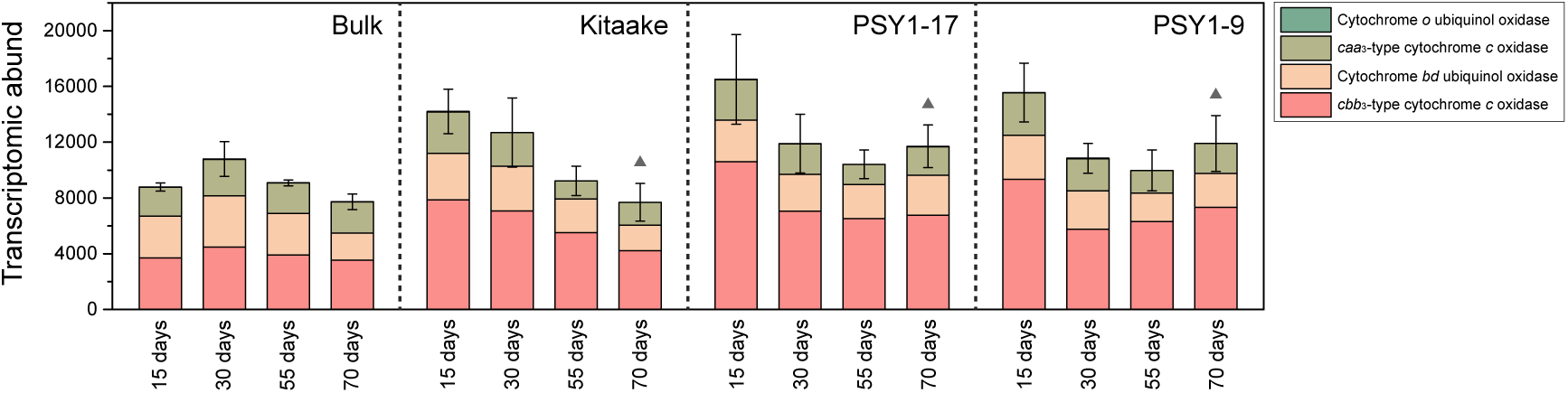
Transcriptomic abundance and category of terminal oxidase genes. Error bars indicate standard deviations of sum values in replicates. Symbols (i.e., triangles) indicate significant differences between the Kitaake and PSY1 genotypes (PSY1-17 and PSY1-9) at the designated developmental stage.

**Supplementary Fig. 4.**
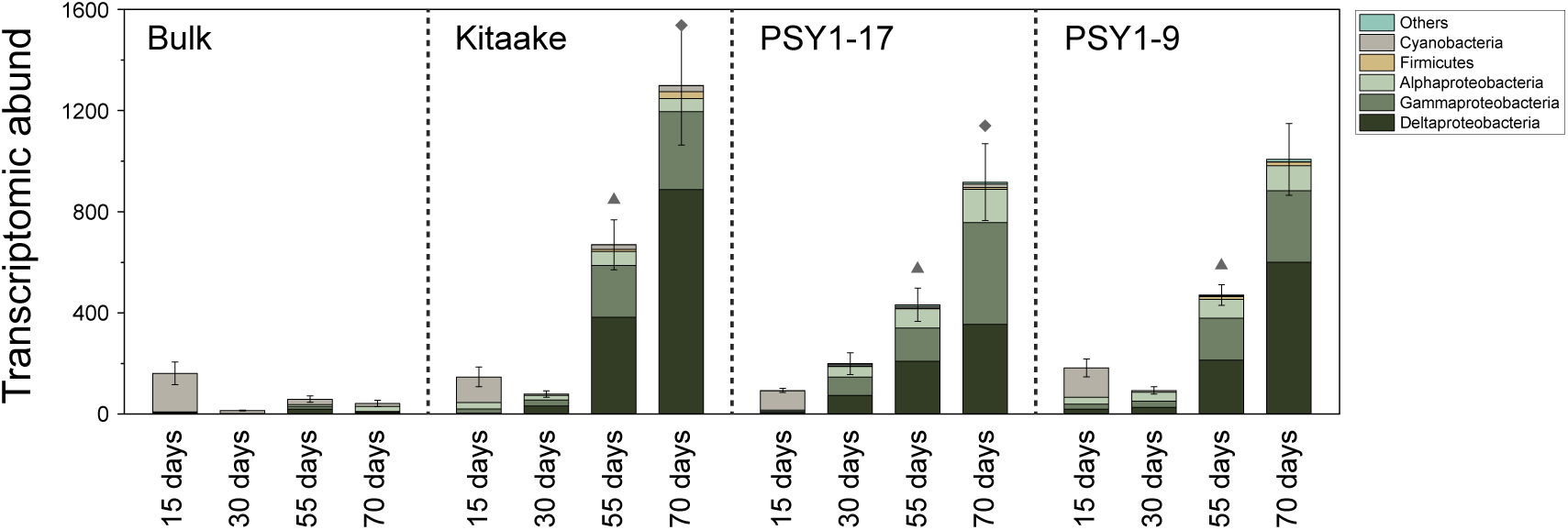
Transcriptomic abundance and taxonomic affiliation of nitrogenase genes. Error bars indicate standard deviations of sum values in replicates. Symbols (i.e., triangles and diamonds) indicate significant differences between Kitaake and PSY1 genotypes (PSY1-17 and/or PSY1-9) at the same developmental stage.

**Supplementary Fig. 5.**
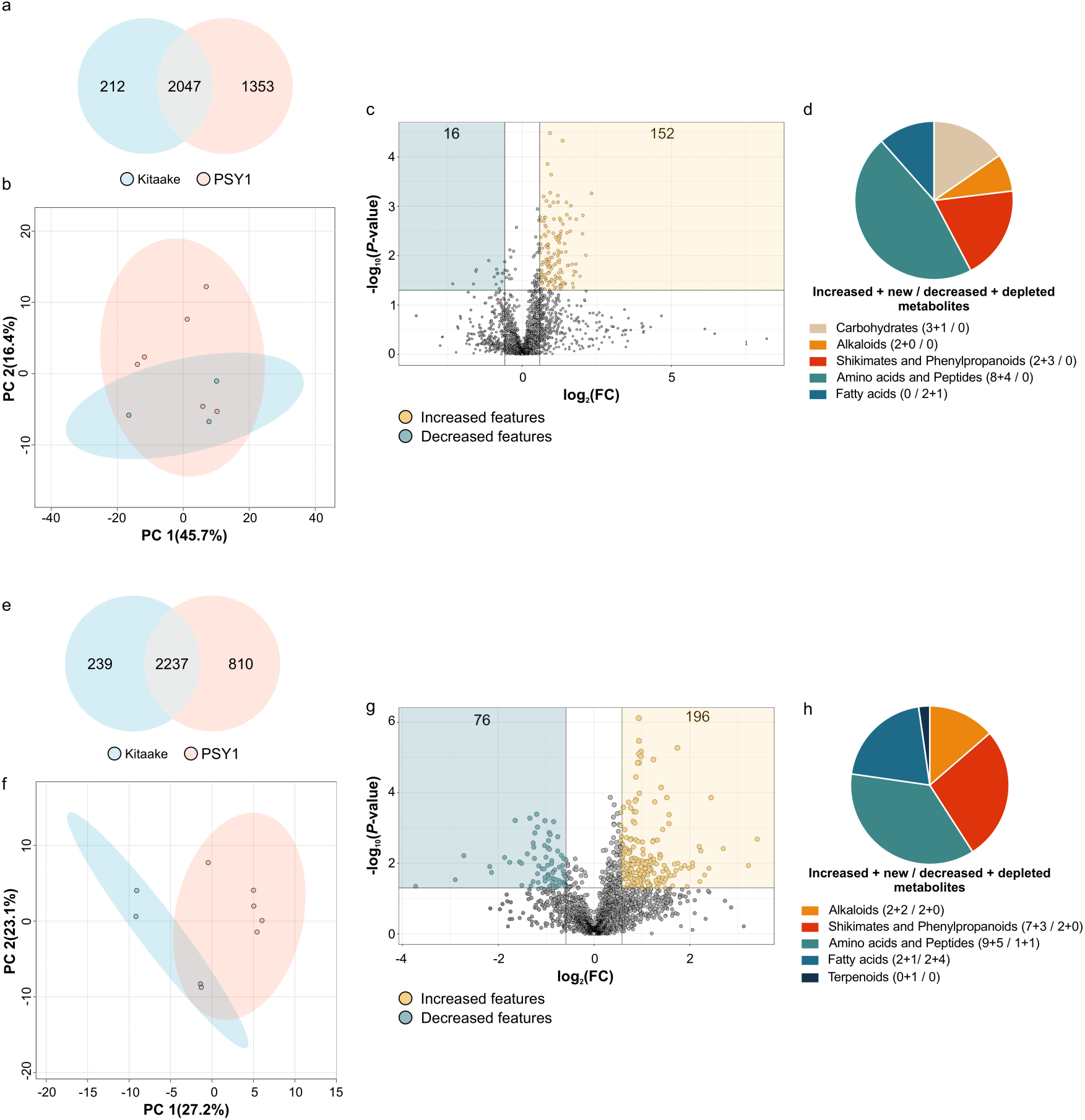
Metabolomic analysis of Kitaake and transgenic PSY1 lines using hydrophilic interaction liquid chromatography (positive ionization mode). **(a** and **e)** Venn diagram of features detected in Kitaake and *Ubi::OsPSY1* (PSY1) samples. (**b** and **f**) Principal component analysis plot (PCA) and (**c** and **g**) volcano plots of features detected in exudates (**a-b**) or tissue (**e-f**) from Kitaake and PSY1 roots, respectively. The number of decreased (in blue) and increased (in yellow) features in transgenic lines compared with Kitaake control is indicated on each plot (log2 fold change +0.5/−0.5 and P < 0.05). Grey dots represent features not differentially abundant. (**d** and **h**) Classification of differentially abundant metabolites identified for root exudates or tissue. A breakdown of increased, new, decreased, and depleted metabolites is indicated next to each color symbol.

**Supplementary Fig. 6.**
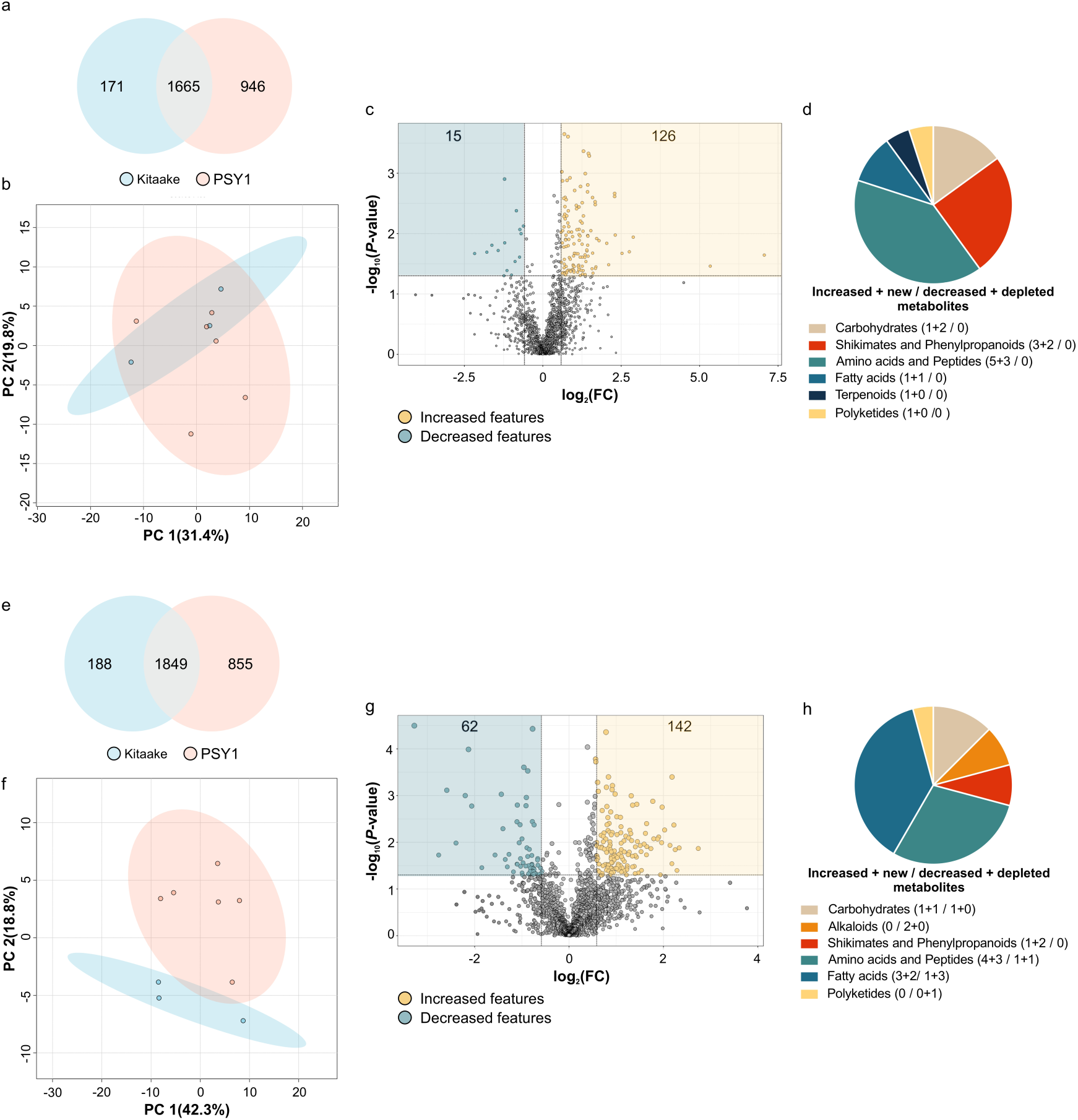
Metabolomic analysis of Kitaake and transgenic PSY1 lines using hydrophilic interaction liquid chromatography (negative ionization mode). **(a** and **e)** Venn diagram of features detected in Kitaake and *Ubi::OsPSY1* (PSY1) samples. (**b** and **f**) Principal component analysis plot (PCA) and (**c** and **g**) volcano plots of features detected in exudates (**a-b**) or tissue (**e-f**) from Kitaake and PSY1 roots, respectively. The number of decreased (in blue) and increased (in yellow) features in transgenic lines compared with Kitaake control is indicated on each plot (log2 fold change +0.5/−0.5 and P < 0.05). Grey dots represent features not differentially abundant. (**d** and **h**) Classification of differentially abundant metabolites identified for root exudates or tissue. A breakdown of increased, new, decreased, and depleted metabolites is indicated next to each color symbol.

**Supplementary Fig. 7.**
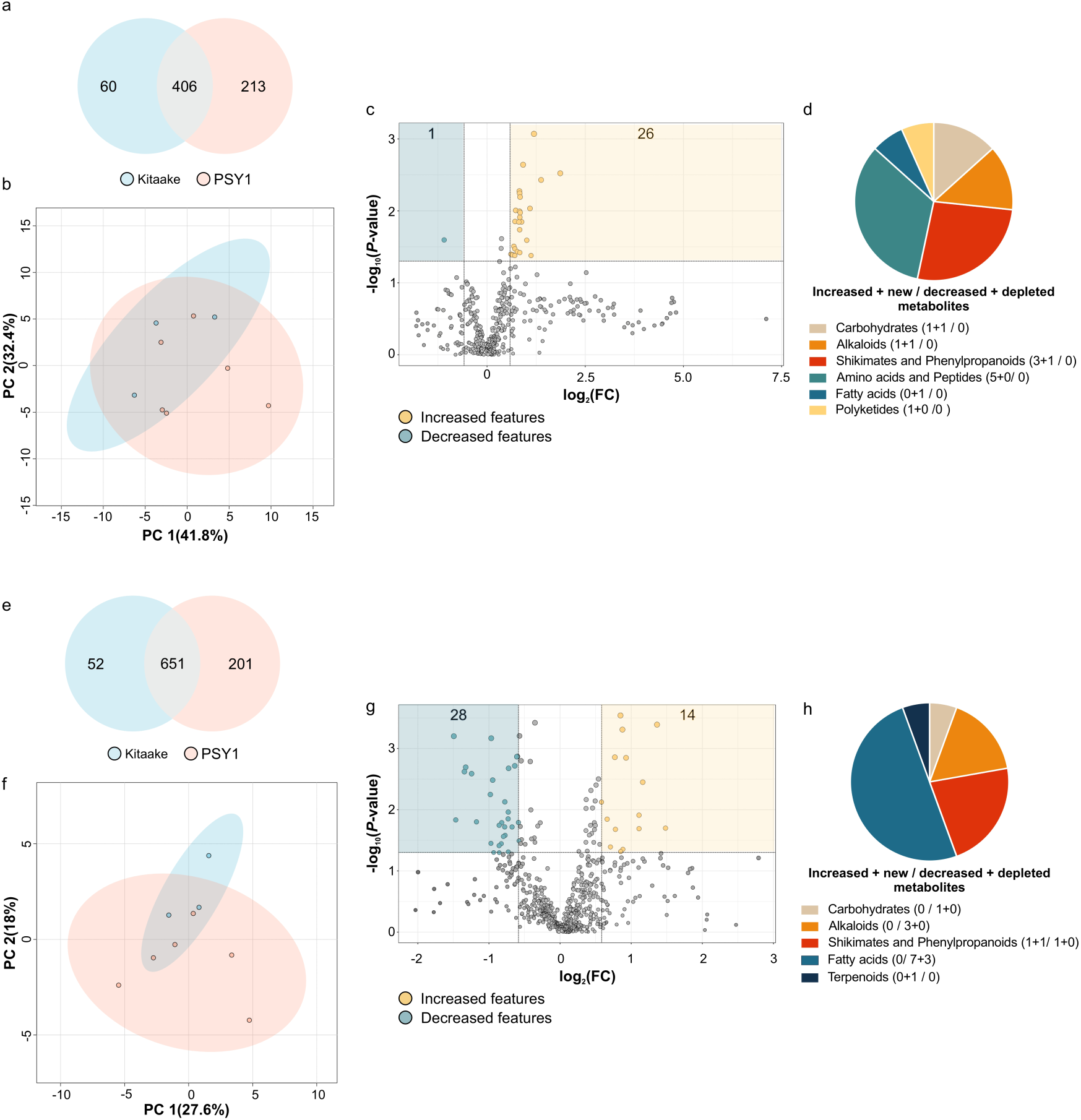
Metabolomic analysis of Kitaake and transgenic PSY1 lines using reverse phase chromatography (positive ionization mode). **(a** and **e)** Venn diagram of features detected in Kitaake and *Ubi::OsPSY1* (PSY1) samples. (**b** and **f**) Principal component analysis plot (PCA) and (**c** and **g**) volcano plots of features detected in exudates (**a-b**) or tissue (**e-f**) from Kitaake and PSY1 roots, respectively. The number of decreased (in blue) and increased (in yellow) features in transgenic lines compared with Kitaake control is indicated on each plot (log2 fold change +0.5/−0.5 and P < 0.05). Grey dots represent features not differentially abundant. (**d** and **h**) Classification of differentially abundant metabolites identified for root exudates or tissue. A breakdown of increased, new, decreased, and depleted metabolites is indicated next to each color symbol.

**Supplementary Fig. 8.**
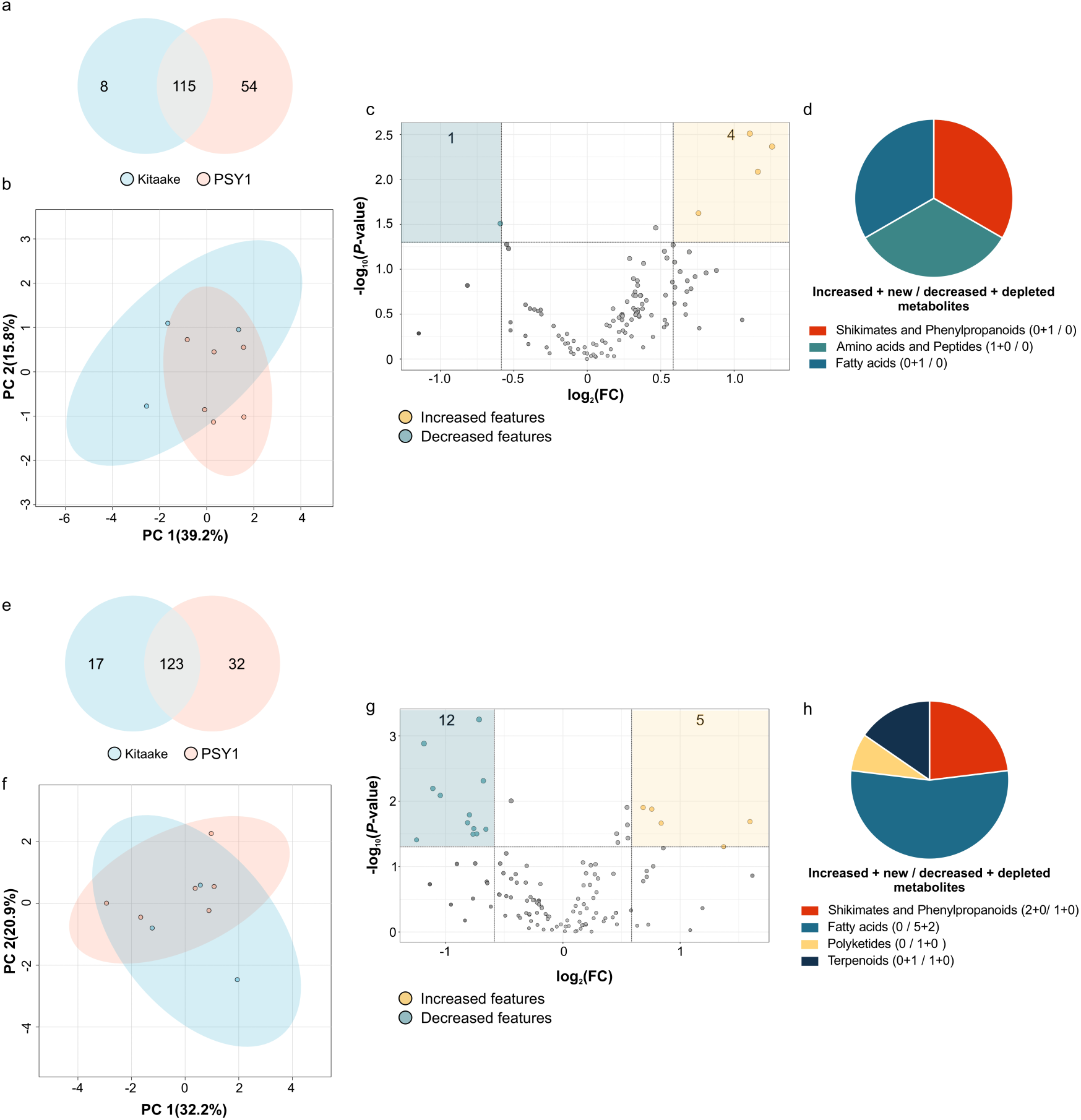
Metabolomic analysis of Kitaake and transgenic PSY1 lines using reverse phase chromatography (negative ionization mode). **(a** and **e)** Venn diagram of features detected in Kitaake and *Ubi::OsPSY1* (PSY1) samples. (**b** and **f**) Principal component analysis plot (PCA) and (**c** and **g**) volcano plots of features detected in exudates (**a-b**) or tissue (**e-f**) from Kitaake and PSY1 roots, respectively. The number of decreased (in blue) and increased (in yellow) features in transgenic lines compared with Kitaake control is indicated on each plot (log2 fold change +0.5/−0.5 and P < 0.05). Grey dots represent features not differentially abundant. (**d** and **h**) Classification of differentially abundant metabolites identified for root exudates or tissue. A breakdown of increased, new, decreased, and depleted metabolites is indicated next to each color symbol.

**Supplementary Fig. 9.**
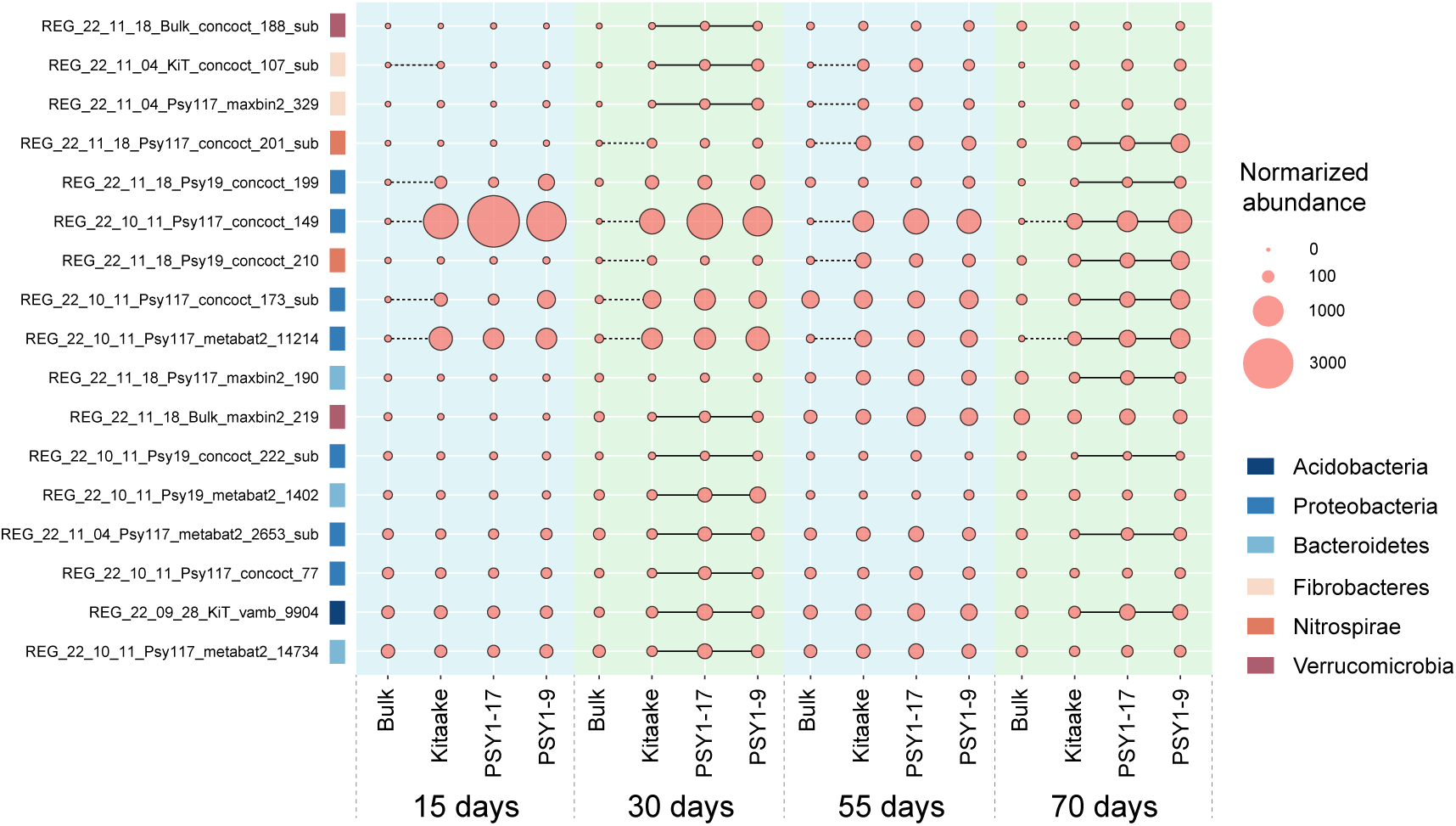
Genomes with activities that are up-regulated in rice PSY transgenic plants. Solid lines linking circles indicate genomes with significantly higher activities in rice mutants than in the wild type (log2FoldChange ≥ 1, P ≤ 0.05), as identified by DESeq2. Dashed lines show that genomes had significantly higher activities in rice planted groups than in bulk soils.

**Supplementary Fig. 10.**
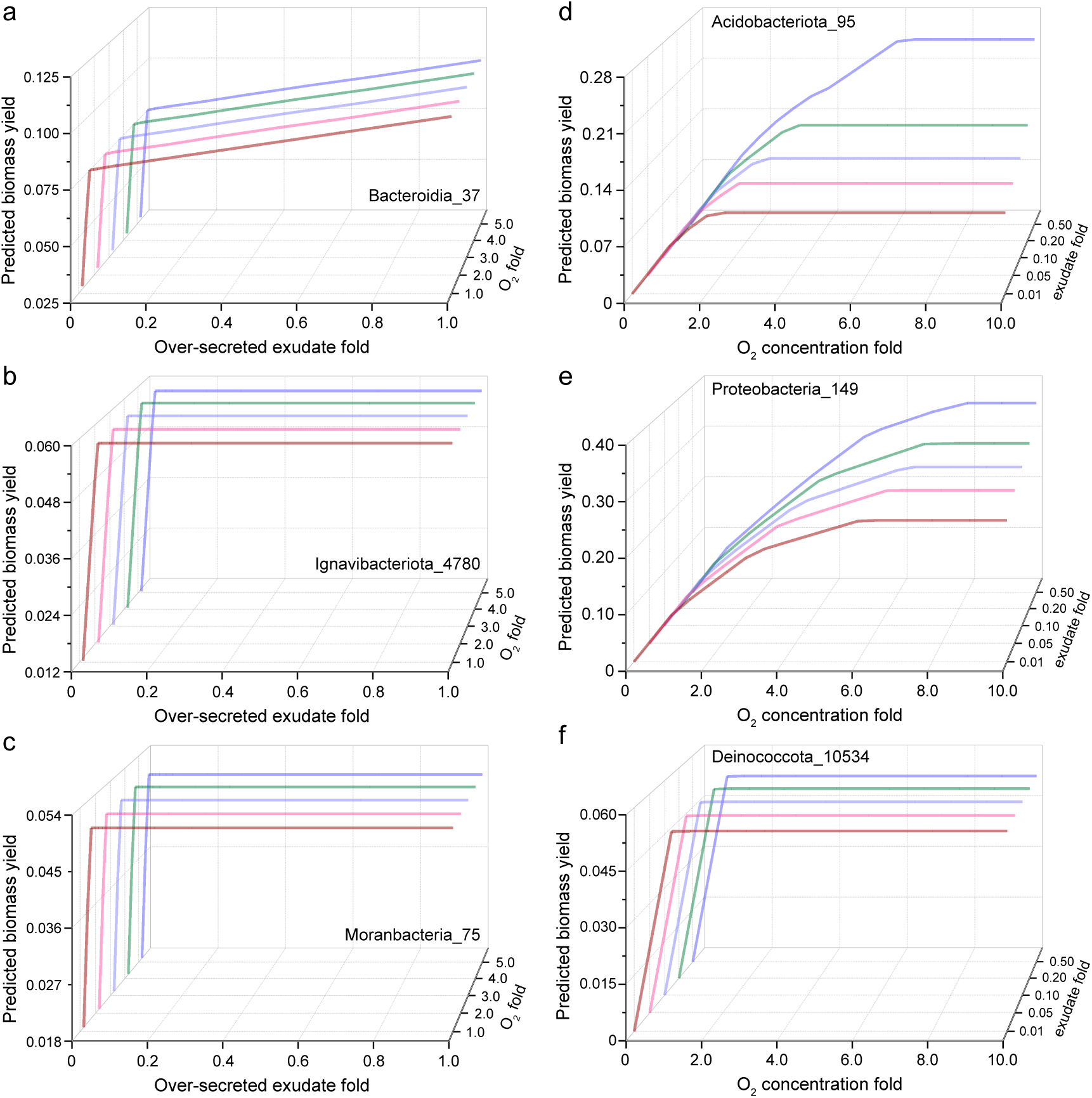
Corresponding biomass yield inferred from metabolic modeling under varying conditions. Each panel has the same setting as the counterpart in Fig. 7.

**Supplementary Fig. 11.**
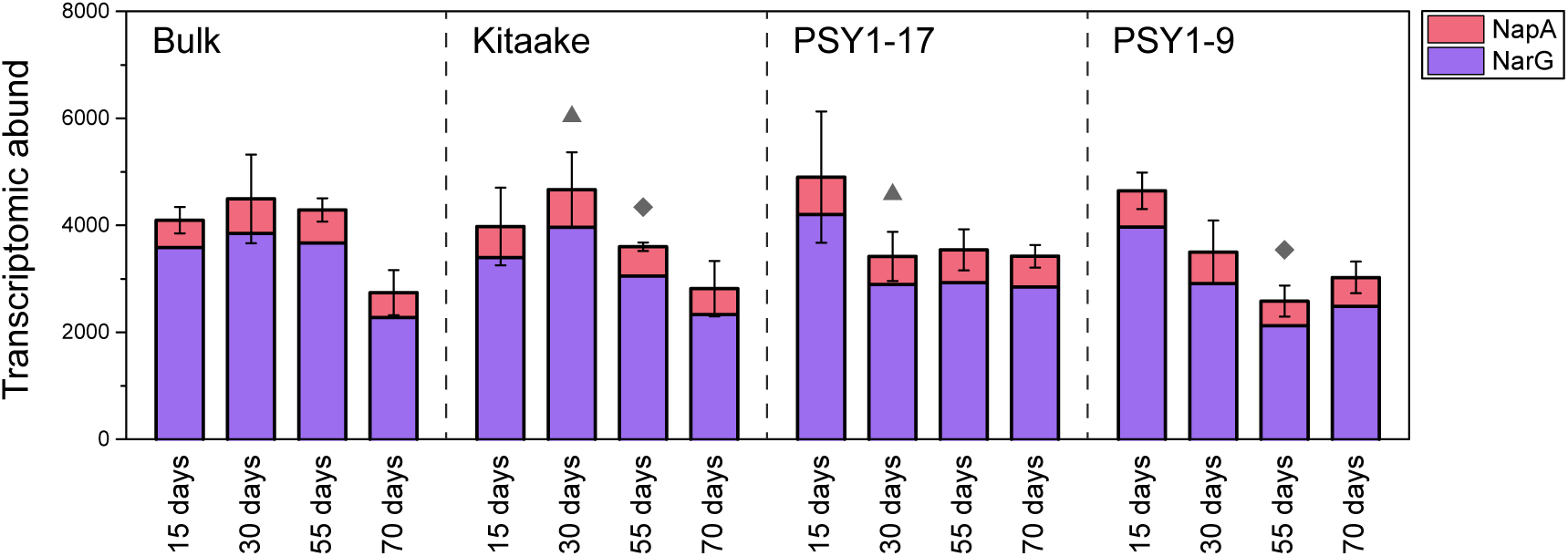
Transcriptomic abundance of nitrate reductase genes. Error bars in the abundance profile indicate standard deviations of sum transcriptomic values in replicates. Symbols (i.e., triangles and diamonds) indicate significant differences between Kitaake and PSY1 genotypes (PSY1-17 and/or PSY1-9) at the same developmental stage.

**Supplementary Fig. 12.**
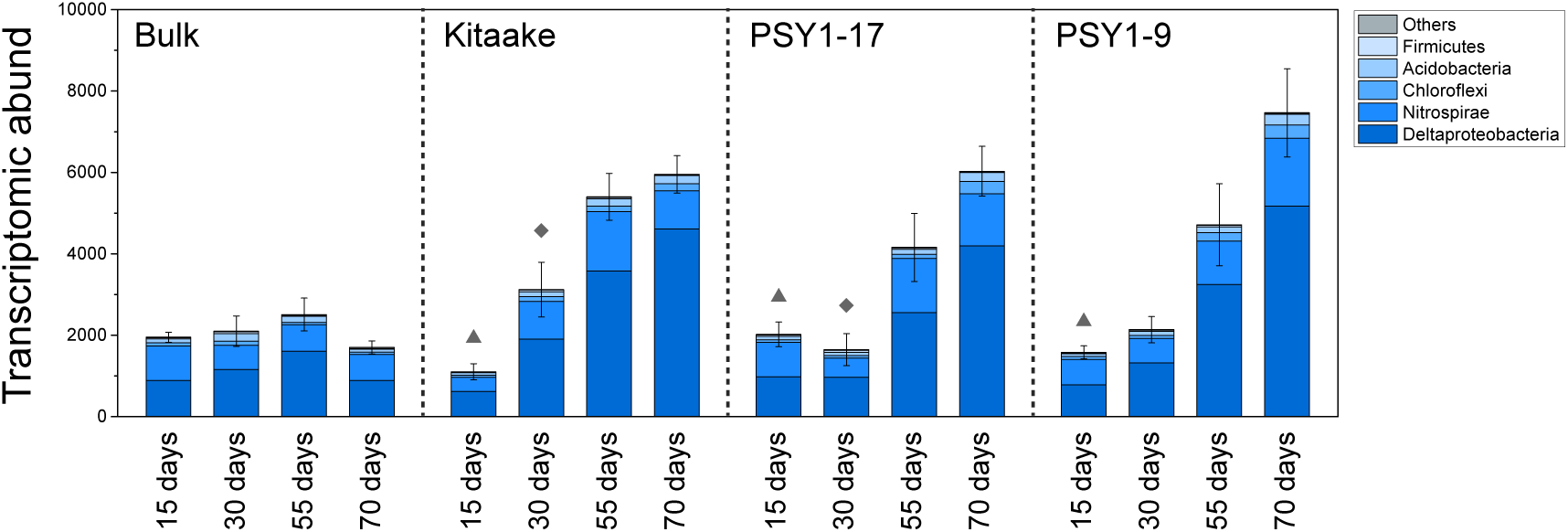
Transcriptomic abundance and taxonomic affiliation of reductive *dsrA*. Error bars indicate standard deviations of sum values in replicates. Symbols (i.e., triangles and diamonds) indicate significant differences between Kitaake and PSY1 genotypes (PSY1-17 and/or PSY1-9) at the same developmental stage.

## References

1. Lan, X., Thoning, K. W. & Dlugokencky E. J. Trends in globally-averaged CH4, N2O, and SF6 determined from NOAA Global Monitoring Laboratory measurements. Version 2024-01, 10.15138/P8XG-AA10. (2022).

2. Liu, Y. L. et al. Rice paddy soils are a quantitatively important carbon store according to a global synthesis. Commun. Earth Environ. 2, (2021).

3. Thauer, R. K., Kaster, A. K., Seedorf, H., Buckel, W. & Hedderich, R. Methanogenic archaea: ecologically relevant differences in energy conservation. Nat. Rev. Microbiol. 6, 579–591 (2008).

4. Raghoebarsing, A. A. et al. A microbial consortium couples anaerobic methane oxidation to denitrification. Nature 440, 918–921 (2006).

5. Cai, Y., Zheng, Y., Bodelier, P. L. E., Conrad, R. & Jia, Z. Conventional methanotrophs are responsible for atmospheric methane oxidation in paddy soils. Nat. Commun. 7, 11728 (2016).

6. Liu, Y. C. & Whitman, W. B. Metabolic, phylogenetic, and ecological diversity of the methanogenic archaea. Incred. Anaerobes Physiol. Genomics Fuels 1125, 171–189 (2008).

7. Bridgham, S. D., Cadillo-Quiroz, H., Keller, J. K. & Zhuang, Q. L. Methane emissions from wetlands: biogeochemical, microbial, and modeling perspectives from local to global scales. Glob. Change Biol. 19, 1325–1346 (2013).

8. Bin Rahman, A. N. M. R. & Zhang, J. H. Trends in rice research: 2030 and beyond. Food Energy Secur. 12, (2023).

9. Qian, H. Y. et al. Greenhouse gas emissions and mitigation in rice agriculture. Nat. Rev. Earth Environ. 4, 716–732 (2023).

10. Su, J. et al. Expression of barley SUSIBA2 transcription factor yields high-starch low-methane rice. Nature 523, 602-+ (2015).

11. Kwon, Y. et al. Loss-of-function gs3 allele decreases methane emission and increases grain yield in rice. *Nat*. Clim. Change 13, 1329–1333 (2023).

12. Yamauchi, T., Abe, F., Tsutsumi, N. & Nakazono, M. Root Cortex Provides a Venue for Gas-Space Formation and Is Essential for Plant Adaptation to Waterlogging. Front. Plant Sci. 10, 259 (2019).

13. Pedersen, O., Sauter, M., Colmer, T. D. & Nakazono, M. Regulation of root adaptive anatomical and morphological traits during low soil oxygen. New Phytol. 229, 42–49 (2021).

14. Chen, Y. et al. Rice root morphological and physiological traits interaction with rhizosphere soil and its effect on methane emissions in paddy fields. Soil Biol. Biochem. 129, 191–200 (2019).

15. Ding, H., Jiang, Y. & Cao, C. Deep rice root systems reduce methane emissions in rice paddies. Plant Soil 468, 337–352 (2021).

16. Amano, Y., Tsubouchi, H., Shinohara, H., Ogawa, M. & Matsubayashi, Y. Tyrosine-sulfated glycopeptide involved in cellular proliferation and expansion in Arabidopsis. Proc. Natl. Acad. Sci. U. S. A. 104, 18333–8 (2007).

17. Pruitt, R. N. et al. A microbially derived tyrosine-sulfated peptide mimics a plant peptide hormone. New Phytol. 215, 725–736 (2017).

18. Ercoli, M. F., et al. An open source plant kinase chemogenomics set. Plant Direct 6, e460 (2022).

19. Ercoli, M. F., Shigenaga, A. M., Teixeira De Araujo, A., Jain, R. & Ronald, P. C. Tyrosine-sulfated peptide hormone induces flavonol biosynthesis to control elongation and differentiation in Arabidopsis primary root. Preprint at 10.1101/2024.02.02.578681 (2024).

20. Aulakh, M. S., Wassmann, R., Bueno, C. & Rennenberg, H. Impact of root exudates of different cultivars and plant development stages of rice (L.) on methane production in a paddy soil. Plant Soil 230, 77–86 (2001).

21. Kesawat, M. S. et al. Genome-wide survey of peptides containing tyrosine sulfation (PSY) gene family and potential PSY specific miRNA revealed their role in plant development and diverse stress conditions in rice (Oryza sativa L.). Plant Stress 11, 100412 (2024).

22. Tost, A. S., Kristensen, A., Olsen, L. I., Axelsen, K. B. & Fuglsang, A. T. The PSY Peptide Family-Expression, Modification and Physiological Implications. Genes Basel 12, (2021).

23. Ogawa-Ohnishi, M. et al. Peptide ligand-mediated trade-off between plant growth and stress response. Science 378, 175–180 (2022).

24. Wang, L. et al. A dynamic gene expression atlas covering the entire life cycle of rice. Plant J. 61, 752–766 (2010).

25. Huang, L. & Schiefelbein, J. Conserved Gene Expression Programs in Developing Roots from Diverse Plants. Plant Cell 27, 2119–2132 (2015).

26. Chandran, A. K. N. et al. Rice Chalky Grain 5 regulates natural variation for grain quality under heat stress. Front. Plant Sci. 13, 1026472 (2022).

27. Counce, P. A., Keisling, T. C. & Mitchell, A. J. A Uniform, Objective, and Adaptive System for Expressing Rice Development. Crop Sci. 40, 436–443 (2000).

28. Sharon, I. et al. Accurate, multi-kb reads resolve complex populations and detect rare microorganisms. Genome Res. 25, 534–43 (2015).

29. Delgado-Baquerizo, M. et al. A global atlas of the dominant bacteria found in soil. Science 359, 320-+ (2018).

30. Ding, L. J. et al. Microbiomes inhabiting rice roots and rhizosphere. Fems Microbiol. Ecol. 95, (2019).

31. Ma, B. et al. A genomic catalogue of soil microbiomes boosts mining of biodiversity and genetic resources. Nat. Commun. 14, 7318 (2023).

32. Cho, S. R. et al. A new approach to suppress methane emissions from rice cropping systems using ethephon. Sci. Total Environ. 804, (2022).

33. Zecchin, S. et al. Microbial communities in paddy soils: differences in abundance and functionality between rhizosphere and pore water, the influence of different soil organic carbon, sulfate fertilization and cultivation time, and contribution to arsenic mobility and speciation. Fems Microbiol. Ecol. 99, (2023).

34. Breidenbach, B., Pump, J. & Dumont, M. G. Microbial Community Structure in the Rhizosphere of Rice Plants. Front. Microbiol. 6, (2016).

35. Grosse, S. et al. Purification and characterization of the soluble methane monooxygenase of the type II methanotrophic bacterium sp strain WI 14. Appl. Environ. Microbiol. 65, 3929–3935 (1999).

36. Garcia, P. S., Gribaldo, S. & Borrel, G. Diversity and Evolution of Methane-Related Pathways in Archaea. Annu. Rev. Microbiol. 76, 727–755 (2022).

37. Lubitz, W., Ogata, H., Rudiger, O. & Reijerse, E. Hydrogenases. Chem. Rev. 114, 4081–4148 (2014).

38. Sondergaard, D., Pedersen, C. N. S. & Greening, C. HydDB: A web tool for hydrogenase classification and analysis. Sci. Rep. 6, 34212 (2016).

39. Greening, C. et al. Genomic and metagenomic surveys of hydrogenase distribution indicate H is a widely utilised energy source for microbial growth and survival. Isme J. 10, 761–777 (2016).

40. Zehr, J. P. & Capone, D. G. Changing perspectives in marine nitrogen fixation. Science 368, 729-+ (2020).

41. Dong, X. Y. et al. Phylogenetically and catabolically diverse diazotrophs reside in deep-sea cold seep sediments. Nat. Commun. 13, (2022).

42. Gralka, M., Pollak, S. & Cordero, O. X. Genome content predicts the carbon catabolic preferences of heterotrophic bacteria. Nat Microbiol 8, 1799–1808 (2023).

43. OECD & Food and Agriculture Organization of the United Nations. OECD-FAO Agricultural Outlook 2021-2030. (OECD, 2021). doi:10.1787/19428846-en.

44. Wang, X. et al. Global and regional trends in greenhouse gas emissions from rice production, trade, and consumption. Environ. Impact Assess. Rev. 101, 107141 (2023).

45. Conrad, R., Klose, M., Lu, Y. & Chidthaisong, A. Methanogenic Pathway and Archaeal Communities in Three Different Anoxic Soils Amended with Rice Straw and Maize Straw. Front. Microbiol. 3, (2012).

46. Hu, J. et al. A low-methane rice with high-yield potential realized via optimized carbon partitioning. Sci. Total Environ. 920, 170980 (2024).

47. Chidthaisong, A. & Conrad, R. Turnover of glucose and acetate coupled to reduction of nitrate, ferric iron and sulfate and to methanogenesis in anoxic rice field soil. Fems Microbiol. Ecol. 31, 73–86 (2000).

48. Krüger, M., Frenzel, P. & Conrad, R. Microbial processes influencing methane emission from rice fields. Glob. Change Biol. 7, 49–63 (2001).

49. Jetten, M. S. M., Stams, A. J. M. & Zehnder, A. J. B. Methanogenesis from Acetate - a Comparison of the Acetate Metabolism in Methanothrix-Soehngenii and Methanosarcina Spp. Fems Microbiol. Lett. 88, 181–197 (1992).

50. Chin, K. J., Lueders, T., Friedrich, M. W., Klose, M. & Conrad, R. Archaeal community structure and pathway of methane formation on rice roots. Microb. Ecol. 47, 59–67 (2004).

51. Lovley, D. R. Minimum threshold for hydrogen metabolism in methanogenic bacteria. Appl. Environ. Microbiol. 49, 1530–1 (1985).

52. Feldewert, C., Lang, K. & Brune, A. The hydrogen threshold of obligately methyl-reducing methanogens. Fems Microbiol. Lett. 367, (2020).

53. Lyu, Z. & Lu, Y. Metabolic shift at the class level sheds light on adaptation of methanogens to oxidative environments. ISME J. 12, 411–423 (2018).

54. Angel, R., Claus, P. & Conrad, R. Methanogenic archaea are globally ubiquitous in aerated soils and become active under wet anoxic conditions. ISME J. 6, 847–862 (2012).

55. Cabrol, L. et al. Microbial ecology of fermentative hydrogen producing bioprocesses: useful insights for driving the ecosystem function. Fems Microbiol. Rev. 41, 158–181 (2017).

56. Kerdchoechuen, O. Methane emission in four rice varieties as related to sugars and organic acids of roots and root exudates and biomass yield. Agric. Ecosyst. Environ. 108, 155–163 (2005).

57. Qian, P. et al. The CLE9/10 secretory peptide regulates stomatal and vascular development through distinct receptors. Nat. Plants 4, 1071–1081 (2018).

58. Zampieri, M., Hörl, M., Hotz, F., Müller, N. F. & Sauer, U. Regulatory mechanisms underlying coordination of amino acid and glucose catabolism in Escherichia coli. Nat. Commun. 10, 3354 (2019).

59. Chandel, N. S. Amino Acid Metabolism. Cold Spring Harb. Perspect. Biol. 13, a040584 (2021).

60. Moe, L. A. Amino acids in the rhizosphere: from plants to microbes. Am J Bot 100, 1692–705 (2013).

61. Aulakh, M. S., Wassmann, R., Bueno, C., Kreuzwieser, J. & Rennenberg, H. Characterization of Root Exudates at Different Growth Stages of Ten Rice (*Oryza sativa* L.) Cultivars. Plant Biol. 3, 139–148 (2008).

62. Schindelin, J. et al. Fiji: an open-source platform for biological-image analysis. Nat. Methods 9, 676–682 (2012).

63. Karimi, M., Depicker, A. & Hilson, P. Recombinational Cloning with Plant Gateway Vectors. Plant Physiol. 145, 1144–1154 (2007).

64. Debernardi, J. M. et al. A GRF–GIF chimeric protein improves the regeneration efficiency of transgenic plants. Nat. Biotechnol. 38, 1274–1279 (2020).

65. Hellemans, J., Mortier, G., De Paepe, A., Speleman, F. & Vandesompele, J. qBase relative quantification framework and software for management and automated analysis of real-time quantitative PCR data. Genome Biol. 8, R19 (2007).

66. Taylor, I., et al. Mechanism and function of root circumnutation. Proc. Natl. Acad. Sci. 118, e2018940118 (2021).

67. Ursache, R., Andersen, T. G., Marhavý, P. & Geldner, N. A protocol for combining fluorescent proteins with histological stains for diverse cell wall components. Plant J. 93, 399–412 (2018).

68. Bruce, R. J. & West, C. A. Elicitation of Lignin Biosynthesis and Isoperoxidase Activity by Pectic Fragments in Suspension Cultures of Castor Bean. Plant Physiol. 91, 889–897 (1989).

69. Nurk, S., Meleshko, D., Korobeynikov, A. & Pevzner, P. A. metaSPAdes: a new versatile metagenomic assembler. Genome Res. 27, 824–834 (2017).

70. Kang, D. W. D. et al. MetaBAT 2: an adaptive binning algorithm for robust and efficient genome reconstruction from metagenome assemblies. Peerj 7, e7359 (2019).

71. Alneberg, J. et al. Binning metagenomic contigs by coverage and composition. Nat Methods 11, 1144–1146 (2014).

72. Wu, Y. W., Simmons, B. A. & Singer, S. W. MaxBin 2.0: an automated binning algorithm to recover genomes from multiple metagenomic datasets. Bioinformatics 32, 605–607 (2016).

73. Nissen, J. N. et al. Improved metagenome binning and assembly using deep variational autoencoders. Nat. Biotechnol. 39, 555–560 (2021).

74. Sieber, C. M. K., et al. Recovery of genomes from metagenomes via a dereplication, aggregation and scoring strategy. Nat Microbiol 3, 836-+ (2018).

75. Olm, M. R., Brown, C. T., Brooks, B. & Banfield, J. F. dRep: a tool for fast and accurate genomic comparisons that enables improved genome recovery from metagenomes through de-replication. ISME J. 11, 2864–2868 (2017).

76. Chklovski, A., Parks, D. H., Woodcroft, B. J. & Tyson, G. W. CheckM2: a rapid, scalable and accurate tool for assessing microbial genome quality using machine learning. Nat Methods 20, 1203–1212 (2023).

77. Chaumeil, P. A., Mussig, A. J., Hugenholtz, P. & Parks, D. H. GTDB-Tk v2: memory friendly classification with the genome taxonomy database. Bioinformatics 38, 5315–5316 (2022).

78. Parks, D. H. et al. A complete domain-to-species taxonomy for Bacteria and Archaea. Nat. Biotechnol. 38, 1079–1086 (2020).

79. Kopylova, E., Noe, L. & Touzet, H. SortMeRNA: fast and accurate filtering of ribosomal RNAs in metatranscriptomic data. Bioinformatics 28, 3211–3217 (2012).

80. Bushmanova, E., Antipov, D., Lapidus, A. & Prjibelski, A. D. rnaSPAdes: a de novo transcriptome assembler and its application to RNA-Seq data. Gigascience 8, (2019).

81. Hyatt, D. et al. Prodigal: prokaryotic gene recognition and translation initiation site identification. BMC Bioinformatics 11, (2010).

82. Steinegger, M. & Soding, J. MMseqs2 enables sensitive protein sequence searching for the analysis of massive data sets. Nat. Biotechnol. 35, 1026–1028 (2017).

83. Steinegger, M. & Soding, J. Clustering huge protein sequence sets in linear time. Nat. Commun. 9, 2542 (2018).

84. Kanehisa, M., Furumichi, M., Sato, Y., Kawashima, M. & Ishiguro-Watanabe, M. KEGG for taxonomy-based analysis of pathways and genomes. Nucleic Acids Res. (2022) doi:10.1093/nar/gkac963.

85. Bateman, A. et al. UniProt: the Universal Protein Knowledgebase in 2023. Nucleic Acids Res. (2022) doi:10.1093/nar/gkac1052.

86. Edgar, R. C. Search and clustering orders of magnitude faster than BLAST. Bioinformatics 26, 2460–2461 (2010).

87. Drula, E. et al. The carbohydrate-active enzyme database: functions and literature. Nucleic Acids Res. 50, D571–D577 (2022).

88. Zheng, J. et al. dbCAN3: automated carbohydrate-active enzyme and substrate annotation. Nucleic Acids Res. 51, W115–W121 (2023).

89. Finn, R. D. et al. The Pfam protein families database: towards a more sustainable future. Nucleic Acids Res. 44, D279–D285 (2016).

90. Eddy, S. R. Accelerated profile HMM searches. Plos Comput. Biol. 7, e1002195 (2011).

91. Olm, M. R. et al. Consistent Metagenome-Derived Metrics Verify and Delineate Bacterial Species Boundaries. Msystems 5, e00731–19 (2020).

92. Rognes, T., Flouri, T., Nichols, B., Quince, C. & Mahe, F. VSEARCH: a versatile open source tool for metagenomics. Peerj 4, (2016).

93. Boratyn, G. M. et al. BLAST: a more efficient report with usability improvements. Nucleic Acids Res. 41, W29–W33 (2013).

94. Katoh, K., Misawa, K., Kuma, K. & Miyata, T. MAFFT: a novel method for rapid multiple sequence alignment based on fast Fourier transform. Nucleic Acids Res. 30, 3059–3066 (2002).

95. Capella-Gutierrez, S., Silla-Martinez, J. M. & Gabaldon, T. trimAl: a tool for automated alignment trimming in large-scale phylogenetic analyses. Bioinformatics 25, 1972–1973 (2009).

96. Nguyen, L. T., Schmidt, H. A., von Haeseler, A. & Minh, B. Q. IQ-TREE: A Fast and Effective Stochastic Algorithm for Estimating Maximum-Likelihood Phylogenies. Mol. Biol. Evol. 32, 268–274 (2015).

97. Letunic, I. & Bork, P. Interactive Tree Of Life (iTOL) v5: an online tool for phylogenetic tree display and annotation. Nucleic Acids Res. 49, W293–W296 (2021).

98. Neukirchen, S. & Sousa, F. L. DiSCo: a sequence-based type-specific predictor of Dsr-dependent dissimilatory sulphur metabolism in microbial data. *Microb*. Genomics 7, (2021).

99. Langmead, B. & Salzberg, S. L. Fast gapped-read alignment with Bowtie 2. Nat Methods 9, 357–U54 (2012).

100. Liao, Y., Smyth, G. K. & Shi, W. featureCounts: an efficient general purpose program for assigning sequence reads to genomic features. Bioinformatics 30, 923–30 (2014).

101. Love, M. I., Huber, W. & Anders, S. Moderated estimation of fold change and dispersion for RNA-seq data with DESeq2. Genome Biol. 15, (2014).

102. Benjamini, Y. & Hochberg, Y. Controlling the False Discovery Rate - a Practical and Powerful Approach to Multiple Testing. J. R. Stat. Soc. Ser. B-Stat. Methodol. 57, 289– 300 (1995).

103. Nothias, L.-F. et al. Feature-based molecular networking in the GNPS analysis environment. Nat. Methods 17, 905–908 (2020).

104. Schmid, R. et al. Integrative analysis of multimodal mass spectrometry data in MZmine 3. Nat. Biotechnol. 41, 447–449 (2023).

105. Wang, M. et al. Sharing and community curation of mass spectrometry data with Global Natural Products Social Molecular Networking. Nat. Biotechnol. 34, 828–837 (2016).

106. Kim, H. W. et al. NPClassifier: A Deep Neural Network-Based Structural Classification Tool for Natural Products. J. Nat. Prod. 84, 2795–2807 (2021).

107. Sumner, L. W. et al. Proposed minimum reporting standards for chemical analysis: Chemical Analysis Working Group (CAWG) Metabolomics Standards Initiative (MSI). Metabolomics 3, 211–221 (2007).

108. Arkin, A. P. et al. KBase: The United States Department of Energy Systems Biology Knowledgebase. Nat. Biotechnol. 36, 566–569 (2018).

109. Dukovski, I. et al. A metabolic modeling platform for the computation of microbial ecosystems in time and space (COMETS). Nat. Protoc. 16, 5030–5082 (2021).

110. Bacilio-Jiménez, M. et al. Chemical characterization of root exudates from rice (Oryza sativa) and their effects on the chemotactic response of endophytic bacteria. Plant Soil 249, 271–277 (2003).

111. Tawaraya, K., Horie, R., Wagatsuma, T., Saito, K. & Oikawa, A. Metabolite profiling of shoot extract, root extract, and root exudate of rice under nitrogen and phosphorus deficiency. Soil Sci. Plant Nutr. 64, 312–322 (2018).

112. Oksanen, J., et al. vegan: Community Ecology Package. R package version 2.6-4. HttpsCRANR-Proj. (2022).

113. Wickham, H. ggplot2: Elegant Graphics for Data Analysis. Springer-Verl. N. Y. (2016).

114. R Core Team. R: A language and environment for statistical computing. R Foundation for Statistical Computing, Vienna, Austria. https://www.R-project.org/. (2022).

115. Leys, C., Ley, C., Klein, O., Bernard, P. & Licata, L. Detecting outliers: Do not use standard deviation around the mean, use absolute deviation around the median. J. Exp. Soc. Psychol. 49, 764–766 (2013).

